# *merlin* v4.0: an updated platform for the reconstruction of high-quality genome-scale metabolic models

**DOI:** 10.1101/2021.02.24.432752

**Authors:** João Capela, Davide Lagoa, Ruben Rodrigues, Emanuel Cunha, Fernando Cruz, Ana Barbosa, José Bastos, Diogo Lima, Eugénio C. Ferreira, Miguel Rocha, Oscar Dias

**Affiliations:** Centre of Biological Engineering, University of Minho, 4710-057, Braga, Portugal; LABBELS – Associate Laboratory, Braga/Guimarães, Portugal

## Abstract

Genome-scale metabolic models have been recognised as useful tools for better understanding living organisms’ metabolism. *merlin* (https://www.merlin-sysbio.org/) is an open-source and user-friendly resource that hastens the models’ reconstruction process, conjugating manual and automatic procedures, while leveraging the user’s expertise with a curation-oriented graphical interface. An updated and redesigned version of *merlin* is herein presented. Since 2015, several features have been implemented in *merlin*, along with deep changes in the software architecture, operational flow, and graphical interface. The current version (4.0) includes the implementation of novel algorithms and third-party tools for genome functional annotation, draft assembly, model refinement, and curation. Such updates increased the user base, resulting in multiple published works, including genome metabolic (re-)annotations and model reconstructions of multiple (lower and higher) eukaryotes and prokaryotes. *merlin* version 4.0 is the only tool able to perform template based and non-template based draft reconstructions, while achieving competitive performance compared to state-of-the art tools both for well and less-studied organisms.

## INTRODUCTION

Genome-scale Metabolic Models (GSMM) are genome-wide representations of a given organism’s metabolism. Accordingly, the metabolic information inferred from the genome is integrated with biochemical data, commonly retrieved from reference databases. Within their broad spectrum of applications (1), high-quality GSMMs can be used to predict phenotypes under different genetic and environmental conditions. Such models have been guiding metabolic engineering towards maximising cell factories’ efficiency, predicting the most suitable conditions for driving flux into the production of compounds of interest.

Over the past two decades, an increasing number of genome sequences have become available (2). Correspondingly, the production of curated GSMMs has taken the pace, achieving the mark of six thousand since 1999’s *Haemophilus influenzae’s* model publication (1). Nevertheless, reconstructing high-quality curated models is often time-consuming and laborious, as it can take from a few months to years (3).

Given the usefulness of GSMMs, efforts have been made to accelerate extensive and time-consuming tasks of the reconstruction process. State-of-the-art platforms can integrate fully and semi-automatic methods and graphical interfaces to assist the model’s manual curation (4). Automatic methods provide valuable clues and resources to hasten the reconstruction of metabolic networks. Nonetheless, manual curation is increasingly recognised as a relevant undertaking to ensure high-quality GSMMs (3), as often absent or incomplete genome annotations (5) and biochemical data lead to poor representations of the organisms’ metabolism. Therefore, a balance between automatic processes and manual curation is desirable.

*merlin* (6) is a comprehensive open-source platform, regularly updated, initially released in 2010 and first published in 2015, aiming to assist and accelerate the main tasks of reconstructing high-quality GSMMs. Since the last major version, multiple tools for genome functional annotation, draft assembly, model refinement, and validation have been implemented. Over the last years, *merlin’s* user base has grown considerably, resulting in the reconstruction of GSMMs of multiple organisms from all domains of life, ranging from small-sized genome bacteria (7) to complex higher eukaryotes, such as the cork oak tree (8). Moreover, the *Kluyveromyces lactis’* model (9), developed and improved with *merlin*, is recognised as a reference among the yeast community, as it has served as the baseline for both experimental studies on metabolism and regulation towards biotechnological applications (10–16) and to build models of other yeasts (17, 18).

Herein, we present the newest version of *merlin* (version 4.0), which includes deep software architecture changes, database management, and significant updates in existing operations. Furthermore, new valuable features have been developed and integrated into the framework, mainly as *plugins*. The graphical interface suffered profound alterations to enhance user-friendliness and the manual evaluation and curation assistance. Moreover, we compared draft GSMMs reconstructed with state-of-the-art tools and *merlin*’s new version to assess the similarity of the draft models to their respective published curated GSMMs. Likewise, we benchmarked *merlin* version 4.0 features to these tools using peer-reviewed criteria.

## MATERIAL AND METHODS

### Software architecture and overview

*merlin* version 4.0 is implemented on top of AIBench, a Java application framework for scientific software development (19) and, as shown in Figure 1, includes four main functional modules: Genome Functional Annotation, Model Reconstruction, Curation, and Data Import/Export. The software architecture is divided into software dependency layers. Herein, two main layers can be considered: *project* and *plugins* (inner shaded grey and outer layers in Figure 1, respectively).

**Figure 1.**
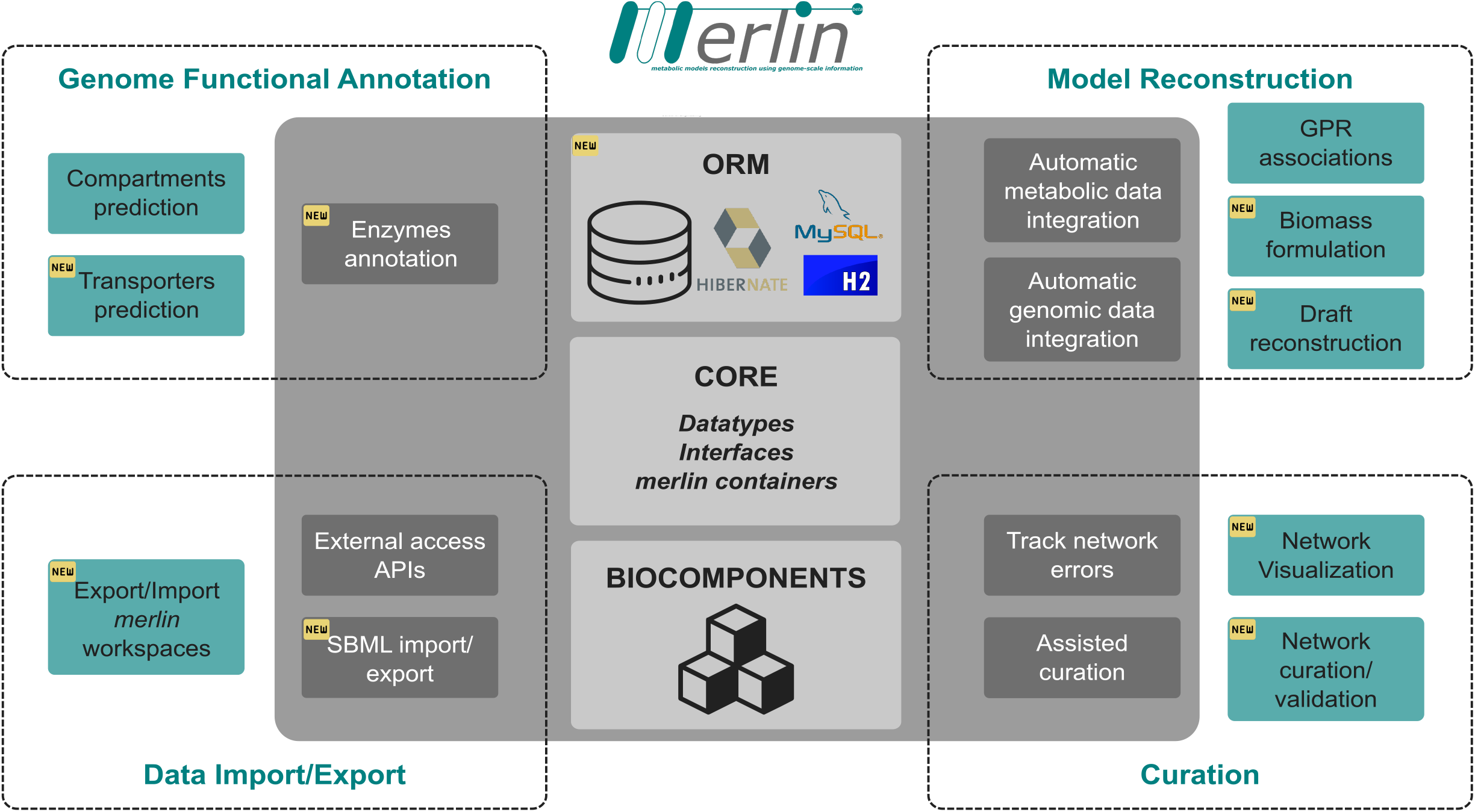
*merlin*’s holistic software architecture. This figure captures the functional modules covered by *merlin* and the layers of software dependencies. The modules are divided into Genome Functional Annotation, Model Reconstruction, Curation, and Data Import/Export. Each of these modules comprises several tools associated with a given reconstruction or data acquisition stage. These tools are either in the *plugins* form (outer, green-shaded boxes) or are included in the software *project* (inner, dark-grey shaded boxes). The *project*, which includes the main components of the software, is also composed of the BIOCOMPONENTS, CORE and ORM modules (central, light-grey boxes). Such components are the cornerstone of the software, as all operations depend on these classes.

The *project*’s *CORE* represents the software’s nuclear layer, where datatypes, interfaces, and containers to manipulate relevant data (e.g., genes, metabolites) are implemented. Moreover, the database access and Object-relational mapping (ORM) modules were integrated into the *project* layer and are presented in Supplementary data 1. The *project*’s *BIOCOMPONENTS* represent the internal library wrappers that allow handling and manipulating a given model without depending on the internal database. Accordingly, these computational objects can be used for independent operations. Finally, *merlin’s project* layer contains the Application Programming Interfaces (APIs) for external data access and other essential operations. Table S1 (Supplementary data 1) provides a brief description of each database used by *merlin* and enumerates the operations that use data retrieved from those sources.

The *plugins’* layer represents optional features that can be both installed and updated at any time. Moreover, a *plugin* management system has been implemented to allow users to update the software, without downloading and installing a new version. Each plugin and its implementation are enumerated and described in Table S2 (Supplementary data 1).

This new architecture is more flexible than the previous, enabling the implementation and release of new features more efficiently.

### Workspaces management

In the previous version, *merlin* allowed deploying several “Projects” simultaneously to reconstruct different GSMMs independently. In *merlin* version 4.0, “Projects” were renamed to “Workspaces”, which better characterises the reconstruction environment. A new feature offered by *merlin* is that “Workspaces” can be exported, imported, and cloned at any moment or stage of the reconstruction process. This helpful feature allows backing up and recovering “Workspaces” at any time or just changing machines without losing any work. Moreover, backward compatibility is guaranteed by a *plugin* that allows importing previous versions of *merlin*’s “Workspaces”.

### Graphical interface updates

*merlin’s* Graphical User Interface (GUI) has significantly changed since the previous published version. Besides alterations in the colours and graphical components, the workspace entities have also changed.

In version 4.0, *model, annotation*, and *validation* are the main modules. Figure S1 (Supplementary data 1) shows that the *model* module is subdivided into five main entities: *genes, proteins, metabolites, reactions*, and *pathways*. The information associated with each of these entities is enumerated in comprehensive tables where users can edit, insert, and remove elements at any moment during the reconstruction process. These entities represent the metabolic information present in the model. The *annotation* module is subdivided into *enzymes* and *compartments*, in which results from the genome functional annotation are enumerated. The *compartments* entity allows users to curate the subcellular localisation prediction results. Lastly, *validation* can include tables with the results retrieved from curation and validation tools, such as *BioISO* (20) and MEMOTE (21).

### New features

*merlin* has several new features on genome’s functional annotations, biomass formulation and network curation, the most relevant of which are presented next.

#### Genome’s functional annotations

*G*enome enzymatic annotation routines in *merlin* include the selection of both gene products and Enzyme Commission (EC) numbers from Basic Local Alignment Search Tool (BLAST) (22) or Diamond (23) alignments’ results. These tools are provided as web applications with APIs that can be accessed externally through *merlin*’s servers. A system of auto-updatable clones of the TrEMBL, Swiss-Prot and RefSeq50 databases, accessible through *merlin*, has been developed to optimise the search for homologous genes.

The scoring algorithm that accounts for the frequency and taxonomy described elsewhere (6, 24) is applied for this selection. In previous versions of *merlin*, the parameterisation of the scoring algorithm was manually and empirically determined, often being a demanding process or not providing the optimal parameters for the genome being annotated. Therefore, *SamPler* (25), a semi-automatic method that relies on statistical metrics and the manual annotation of a genome sample, was developed to improve the optimal parameters’ determination. This tool is available as a *plugin* to configure *merlin*’s annotation routine semi-automatically. Despite being helpful and offering optimum results, *SamPler* may be too demanding for certain purposes. Thus, an automatic procedure is also available for this task. *merlin* provides the *automatic workflow* operation that annotates genes according to a taxonomically related genus or species list. Such a list prioritises the gene product and EC number from entries associated with the selected species or genus. Hence, homologous genes from taxonomically related organisms can be selected as the most suitable candidates for the gene product and EC number annotation.

*TranSyT*, the Transporter Systems Tracker (26), was developed to address the transport systems annotation matter. *TranSyT* uses the Transporter Classification Database (TCDB) (27) as the primary data source for similarity searches and retrieves information on substrates, mechanisms and transport direction. Simultaneously, *TranSyT* uses MetaCyc and the Kyoto Encyclopedia of Genes and Genomes (KEGG) (28) to enrich this information. The transport reactions are created and integrated into the metabolic model automatically.

Model compartmentalisation tools require loading reports from *WolfPSORT* (29), *PSORTb3* (30), or *LocTree3* (31). Compared to previous versions, *merlin* can no longer use a remote Java API for accessing *WolfPSORT*’s functionalities, as it is currently unavailable. Instead, it offers operations to integrate each tool’s prediction report rapidly. For *WolfPSORT* and *LocTree3* reports, *merlin* reads the prediction results directly on the web, requiring only the Uniform Resource Locator (URL) of the report webpage, whereas, for *PSORTb3, merlin* parses a prediction file (“Long Format”) and integrates the results in the database. *LocTree3* reports allow widening the range of options for subcellular localisation prediction of enzymes and metabolites (31). These annotations can be integrated into the model by defining thresholds for the subcellular localisation prediction scores.

#### Biomass formulation

The most common approach towards biomass formulation includes adopting biomass equations from taxonomically related organisms. Although recurrent and assumed not to propagate significant errors (32), this method has been suggested inadequate by a more recent study (33). Instead, estimating the average protein and (deoxy)ribonucleotide contents from the genome seems to provide better predictions (33). Hence, *merlin* provides an operation that calculates the contents, as mentioned above, automatically. Furthermore, gene expression data can be used to adjust the protein contents to experimental data.

Moreover, *merlin* provides templates for creating equations that represent the biomass composition. These templates include the average contents of each biomass-related macromolecule for different types of organisms. These values were retrieved from the literature, the ModelSEED database (34), or experimentally determined. Nevertheless, it is worth noting that these templates should serve as baselines, requiring further curation and adjustments.

#### Network curation

Debugging large networks can be time-consuming, even for the most experienced curator. *merlin*’s graphical interface and services provide the means to hasten such a laborious task. The network topology could already be assessed using the “*Draw in Browser*” functionality, which allows visualising coloured KEGG pathways in the default browser. Now, two new plugins - *MetExploreViz* (35) and *Escher Maps* (36) - complement the network topology analysis package, allowing visualising more than one pathway simultaneously, and highlighting network characteristics such as compartments, shared pathways, and network gaps.

Although these tools ease manual curation, they cannot evaluate which reactions are carrying flux, which metabolites are not being produced, nor assess the model’s consistency. Therefore, new *plugins* have been developed, the Biological networks In Silico Optimization (*BioISO*) and MEMOTE. *BioISO* highlights whether a set of reactions carries flux when maximised or minimised. Also, it enables tracking errors that impair the synthesis of a given reaction’s metabolites. Together with *merlin*’s user-friendly graphical interface (which allows adding, editing, and removing reactions), this tool assists in manual curation and gap-filling.

MEMOTE constitutes a suite of standardised tests proposed by the modelling community to assess the model’s quality. The access to the MEMOTE test suite is implemented through an internally developed API, using the Docker image provided by the authors. The results retrieved by the API are parsed and rendered in comprehensive tables. Thus, the reconstructed models’ quality can be verified without leaving *merlin’s* graphical interface.

#### Other features

*merlin* now allows generating a draft reconstruction from other models. This feature uses alignments to determine which reactions to inherit from the input model. The output is a draft reconstruction, ready for refinement and curation. Moreover, a new in-house tool was implemented to generate draft models from the BiGG models’ database. The BiGG Integration Tool (BIT) (37) is implemented within *merlin* as a *plugin* and allows to select BiGG template models to perform the draft reconstruction. BIT runs bidirectional BLAST alignments between the studied organism’s genome and the selected templates. The BiGG reactions of the template models are mapped according to the associated homologous genes. Finally, the gene-protein-reaction (GPR) rules are propagated using the rules described in (37)

*merlin* version 4.0 also allows importing and exporting the genome, annotating results in the GenBank (38) file format, and GSMMs in various levels and versions of the Systems Biology Markup Language (SBML) (39) format (level 2 versions 1 to 4 and level 3 versions 1 and 2) at any stage of the reconstruction.

### Comparison with the previous version

We compared the runtimes of the European Bioinformatics Institute’s (EBI) remote BLAST with the new implementations of BLAST and Diamond in *merlin*’s servers. However, the former operation was integrated with the retrieval and upload of the homologous genes’ information into *merlin*’s database; thus, the total runtime of the operation was quantified. Therefore, we assessed the alignments runtime and the total runtime of the operation to benchmark *merlin*’s new version.

Moreover, the execution time of other operations, such as the transporters annotation and GPR associations, was also computed.

The operations execution time was quantified on a personal computer with an Intel(R) Core(TM) i7-9700 CPU and 6 gigabytes of allocated RAM.

### Comparison with other tools

#### Assessed reconstruction tools

We compared the draft models reconstructed with state-of-the-art tools, including *merlin*, to curated and published GSMMs to assess each tool’s performance. This analysis accounted for the draft metabolic reconstructions of template-based approaches (AuReMe (40), *merlin-BIT* (37), CarveMe (41), ModelSEED (34), and RAVEN (42)) and non-template-based approaches (autoKEGGRec (43), *merlin*, *and* PathwayTools (44)). Hence, draft models of two prokaryotes, namely *Lactobacillus plantarum* WCFS1 and *Bordetella pertussis Tohama I* - gram-positive and gram-negative bacteria, respectively - were reconstructed using these tools. Moreover, draft GSMMs of the eukaryote *Toxoplasma gondii* were generated with all the above-mentioned tools, except CarveMe and ModelSEED, which have not been developed to reconstruct models of eukaryotes (apart from plants for ModelSEED). The reconstruction procedure for each tool is described in detail in Supplementary data 2.Regarding *merlin*, we generated several models using different workflows to validate all the new and updated features. The methodology, and each step of the workflows, are described in detail in Supplementary data 2.

These draft reconstructions were compared with manually curated and validated models—the GSMMs of *L. plantarum* (45)*, B. pertussis* (46), and *T. gondii* (47), respectively. The former two models are in BiGG format and were selected because they were used to benchmark reconstruction tools in a recent systematic assessement (4), while the last is in the KEGG format and was selected to evaluate each tool’s scalability when reconstructing a less studied (up to April 2022 only 100 entries at Swiss-Prot, while *L. plantarum* has 513 *and B. pertussis* has 1,942) and more complex organism. This comparison included the gene and reaction sets within each model.

#### Template models

Several of the benchmarked tools require template models to generate draft GSMMs. Thus, curated GSMMs of taxonomically related organisms were selected. The following BiGG models were used as templates to reconstruct the draft GSMM:

- *Escherichia coli* str. K-12 substr. MG1655 (iML1515) for *B. pertussis* (gram-negative bacteria);
- *Lactococcus lactis* subsp. cremoris MG1363 (iNF517) for *L. plantarum* (gram-positive bacteria);
- *Plasmodium falciparum 3D7* (iAM_Pf480) for *T. gondii* (human parasite eukaryotes).

#### Comparison of reaction sets

The reactions’ identifiers were converted into the reference model reactions’ identifiers format using MetaNetX (48). In this comparison, the transport, exchange, sink and demand reactions were not considered, as the identifiers’ conversion was not trivial for all the models.

The comparison of the reaction sets between the draft and curated models considered the following:

- True Positives (TP) - reactions of the draft model with at least one *alias* in the curated model;
- False Positives (FP) - reactions of the draft model without any *alias* in the curated model;
- False Negatives (FN) – reactions present in the curated model but absent in the draft model.

#### Comparison of gene sets

The evaluation of draft and curated gene sets encompassed a case insensitive comparison of the locus tag present in the models.

The comparison of the gene sets between the draft and curated models considered the following:

- True Positives (TP) - genes present in the draft and curated models;
- False Positives (FP) - genes present in the draft models and absent in the curated model;
- False Negatives (FN) - absent genes in the draft models but present in the curated model.

#### Metrics

Precision, Recall, TP/FP Ratio, F1 and Jaccard Distance were used for this evaluation and calculated as follows:

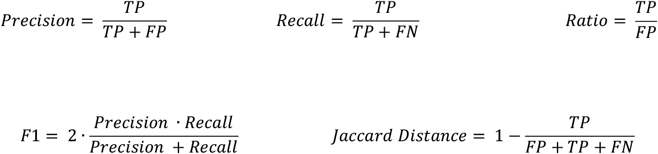

The efforts regarding the curation of draft models are mostly associated with removing FP, which from a curator’s perspective, might be less tedious than adding high volumes of missing information. Thus, high Recall implies that there is not much missing information, as FN are considerably lower than TP. On the other hand, models with higher Precision have a high number of correctly assigned and a low number of incorrectly predicted reactions. Nevertheless, models with high Precision and low Recall have massive volumes of missing information. Accordingly, Recall may be considered a critical metric even in detriment of Precision.

The F1 score combines Precision and Recall into a single metric, allowing a direct evaluation of the similarity of draft and curated models. The Ratio between the reactions correctly inserted (TP) and the additional information (FP) was applied to the draft models. Moreover, the Jaccard Distance (JD) was calculated to assess how different are the draft models from the curated ones. High Ratios and low JDs are desirable when reconstructing draft models.

#### Parameters to assess reconstruction tools

We evaluated the main features of each state-of-the-art tool herein under study. Thus, we evaluated parameters associated with critical steps in the model reconstruction process according to (1, 3, 4, 49, 50): 1) the capacity of performing a genome (re-)annotation; 2) compartmentalisation; 3) generating GPR rules; 4) annotating genes associated with transport reactions; and 5) gap-filling. In addition, features to improve curation were considered, such as: 6) the support to pathway visualisation tools; 7) standard testing with MEMOTE, recognised as a priority for model standardisation by many authors (4, 51). Finally, we assessed other important but general features according to (1, 4, 49) including: 8) compliance with the last version of SBML format; 9) tools to reconstruct a model from scratch (nontemplate-based modelling) or 10) from template models; and 11) the capability of reconstructing GSMMs of eukaryotes. Finally, we also noted 12) whether a GUI was available and if 13) a software license was required to reconstruct a model.

## RESULTS AND DISCUSSION

### Comparison with the previous version

*merlin’s modus operandi* has changed since 2015’s published version. As depicted in Figure 2, the overall reconstruction workflow encompasses four stages, namely 1) Enzymes’ annotation; 2) Draft assembly; 3) Network curation; 4) Model refinement. An enhanced user-friendly interface assists the whole operating flow that leverages the users’ expertise and the curator’s quality.

**Figure 2.**
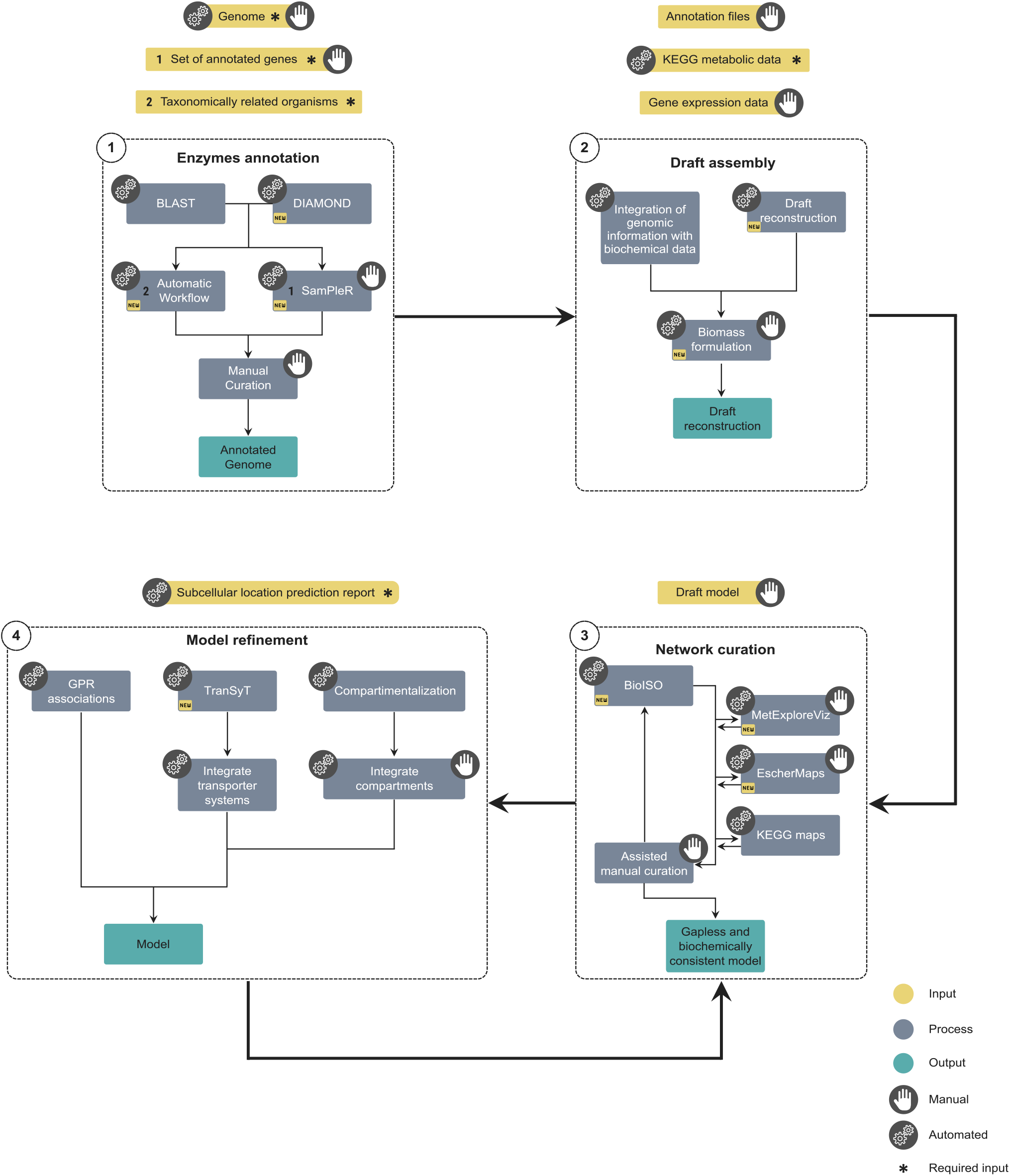
*merlin’s* updated overall reconstruction workflow, including reconstruction steps, possible inputs/outputs, and processes. The “hand” symbol indicates manual processes, while the “gear” symbolises the automatic ones, and both symbols the semi-automatic ones. Each stage’s possible inputs are shown in yellow boxes, processes in blue, and outputs in green. Finally, the asterisk corresponds to the processes’ required inputs. The workflow encompasses four main stages: 1) Enzymes’ annotation; 2) Draft assembly; 3) Model curation; and 4) Model refinement. The output of each stage may not be required for the next, as *merlin* provides operations to import external data. Hence, users can reconstruct models from scratch or external annotations and models.

Before starting a new GSMM reconstruction, users must create a new (or select an existing) *Workspace* using the menu *workspace -> open* at the top bar illustrated in Figure S1. When creating a new Workspace, the user may automatically import an organism’s genome, while, as highlighted in Figure 2, manually importing the genome is also possible.

Although this step is similar to the previous version, the following stages required updating the workflow, which encompasses four steps, as shown in Figure 2 and described next.

#### 1) Enzymes’ annotation

The first step is to automatically annotate the enzymes with either BLAST (22) or Diamond (23) through the *annotation -> enzymes -> BLAST* or *Diamond* operations. The BLAST alignments against Swiss-Prot can take up to 1 hour for about 3000 coding sequences (e.g., *L.plantarum*) and around 6 hours for 8000 coding sequences (e.g., *T. gondii*). Notably, even the slowest Diamond mode (ultra-sensitive) takes only 7 minutes for smaller proteomes and around 13 minutes for longer proteomes. The new features and the speed improvement on the alignments represent significant advances over the previous versions. Loading these high volumes of homologous information, e.g. gene product, EC numbers, and associated organisms, into the user’s database may take about one hour for organisms with larger proteomes. In the previous version, depending on the EBI server traffic and request acceptance rate, these operations (alignments, data retrieval and loading) could take up to 25 hours for the smaller proteomes and 52 hours for larger proteomes. Nevertheless, the alignments continue to be run remotely, now in *merlin*’s servers, without any burden over the user’s personal computer.

Alternatively, annotation reports may be imported to hasten the process of obtaining a final genome annotation through the *annotation -> enzymes -> load* menu. Likewise, existing genome annotations, e.g. KEGG annotations, can be loaded for reannotation using the enhanced GUI.

The next step is applying the scoring system described in (6) to the similarity search results. The best parameters for the re-annotation are selected using a new *plugin, SamPler*, at *annotation -> enzymes -> SamPler*. Alternatively, the scoring may be overruled with the *automatic workflow* at *annotation -> enzymes -> automatic workflow*. The output of these semi- and automatic methods will be the annotation of enzymes with EC numbers. The results will be available at the *enzyme’s* view, as shown in Figure S2 (Supplementary data 3).

Nonetheless, the manual curation of these results is desirable, and such a process is highly facilitated by the graphical interface provided by *merlin*, as shown in Figure S2. The *enzymes* view (Figure S2) provides information about the annotation state, gene products, EC numbers, and scores. Each row’s magnifier button provides information on the BLAST (22) or Diamond (23) operations results. Moreover, the *status* column highlights the enzyme’s revision state, following the pattern described in (24) and redirecting users to the UniProtKB (52) database site. The candidate gene products and EC numbers are available in the dropdown boxes, easily updated and integrated into the model if necessary.

#### 2) Draft assembly

The next stage (Figure 2) is integrating the curated enzymes annotation with metabolic information (metabolites and reactions) retrieved from KEGG. Additionally, finalising the so-called draft metabolic network assembly demands adding pseudo-reactions representing the biomass composition, using the *plugin* at *model -> create -> e-biomass equation*. The biomass pseudo-reactions and KEGG information will be detailed in the *reactions’* (Figure S3, Supplementary data 3), *metabolites’*, and *pathways’* views. This information can be updated at any stage of the reconstruction. Alternatively, a draft model can be automatically reconstructed using the *model -> draft model reconstruction* menu. The draft model can be generated with BiGG metabolic models or previously created workspaces.

#### 3) Network curation

Gaps and inconsistencies are likely to be found in the draft reconstruction. Hence, as highlighted in the third stage of the workflow (Figure 2), *merlin* provides several tools to highlight blocked and unbalanced reactions and correct reactions’ reversibility, among others, to assist in the network manual curation. Additionally, the *reactions* view (Figure S3, Supplementary data 3) allows users to rapidly insert, edit, duplicate and remove existing reactions through explicit buttons and checkboxes. This view also enables the visualisation of reactions from the universal internal database and facilitates their inclusion in the model, easing manual gap-filling.

The new *BioISO* analysis (available at *validation -> BioISO -> execute BioISO*) provides valuable insights into the network’s state. After running *BioISO*, a table is docked in the dashboard’s *validation* module, rendering the results as shown in Figure S4 (Supplementary data 3). Likewise, other tools such as *MetExploreViz, Escher Maps* (accessed through *validation -> network visualisation*), and the *Draw in Browser* operation (Figure S3) can be used to get insights into the network topology.

#### 4) Model refinement

The model compartmentalisation through *PSort3b, LocTree3* or *WolfPSort* is advisable. External reports can be uploaded into *merlin* by executing *annotation -> compartments -> load reports*. The loaded reports’ results will be rendered at the *compartments* view docked in the dashboard’s *enzymes* module, as shown in Figure S5 (Supplementary data 3). Further integration through *annotation -> compartments -> integrate to model* is mandatory, allowing the user to define thresholds over the obtained prediction scores and ignore possible erroneous compartments.

The transporter systems annotation and transport reactions generation can be performed with a novel tool, initially developed for *merlin* but now also available at KBase (53), *TranSyT*, by executing *model -> create -> transport reactions TranSyT*. These reactions will be available in the *reactions* view, associated with a surrogate pathway designated *Transporters Pathway. TranSyT* takes up to four minutes for small-sized networks, such as bacterial, up to eight minutes for larger networks.

Finally, the GPR rules are automatically generated using BLAST or Diamond alignments’ through *merlin’*s operation *model -> create -> gene-protein-reaction associations*. This takes approximately one hour in version 4.0 for around 3000 coding sequences and a metabolic network with nearly 1800 reactions, while in the previous version took around three hours. Note that the time required to perform this step can change depending on KEGG’s server availability.

#### Interoperability and compliance with standards

The MEMOTE test suite can assess the model’s compliance with standards. This *plugin* is available through *validation -> memote*, and the report will be rendered in a comprehensive table, docked under the *validation* module of the dashboard (Figure S6, in Supplementary data 3).

Lastly, the model can be exported (*workspace -> export -> model*) in the SBML format with valid identifiers and cross-references for metabolites and reactions. Other available options are to export in the GenBank or Excel Workbook formats.

*merlin* provides an optimised workflow that allows users to start the reconstruction process to create a high-quality GSMM from scratch. Nevertheless, the workflow’s flexibility, complemented by import/export operations, enables users to begin or continue the reconstruction process from external genome annotations or external GSMMs. This also enables the curation, reannotation, and refinement of already reconstructed models and annotated genomes with *merlin*’s internal tools.

### Comparison with other tools

This study systematically reconstructed 13 draft GSMMs per organism to evaluate gene and reaction contents with different tools and approaches. For each organism, a GSMM was computed per each third-party tool and seven models were reconstructed using *merlin* with different workflows. From the *merlin* options, *merlin-BIT* and *merlin-BS* (using BLAST + *SamPler*) were selected as the best performing. The former is a new template-based approach, while the latter delivers the best nontemplate-based models in most cases. Details on the analysis of the *merlin*-based models are given in Supplementary data 2. Finally, eight draft models - *merlin-BIT, merlin-BS*, AuReMe, CarveMe, autoKEGGRec, ModelSeed, PathwayTools and Raven - were compared to the curated model.

#### Models’ assessment

##### Bordetella pertussis

Regarding the assessment of the models reconstructed for *B. pertussis*, Figure 3 shows all the metrics for reactions (Figure 3A and 3C), and genes (Figure 3B and 3D). The results are detailed in Tables S5 and S6 (Supplementary data 2).

**Figure 3.**
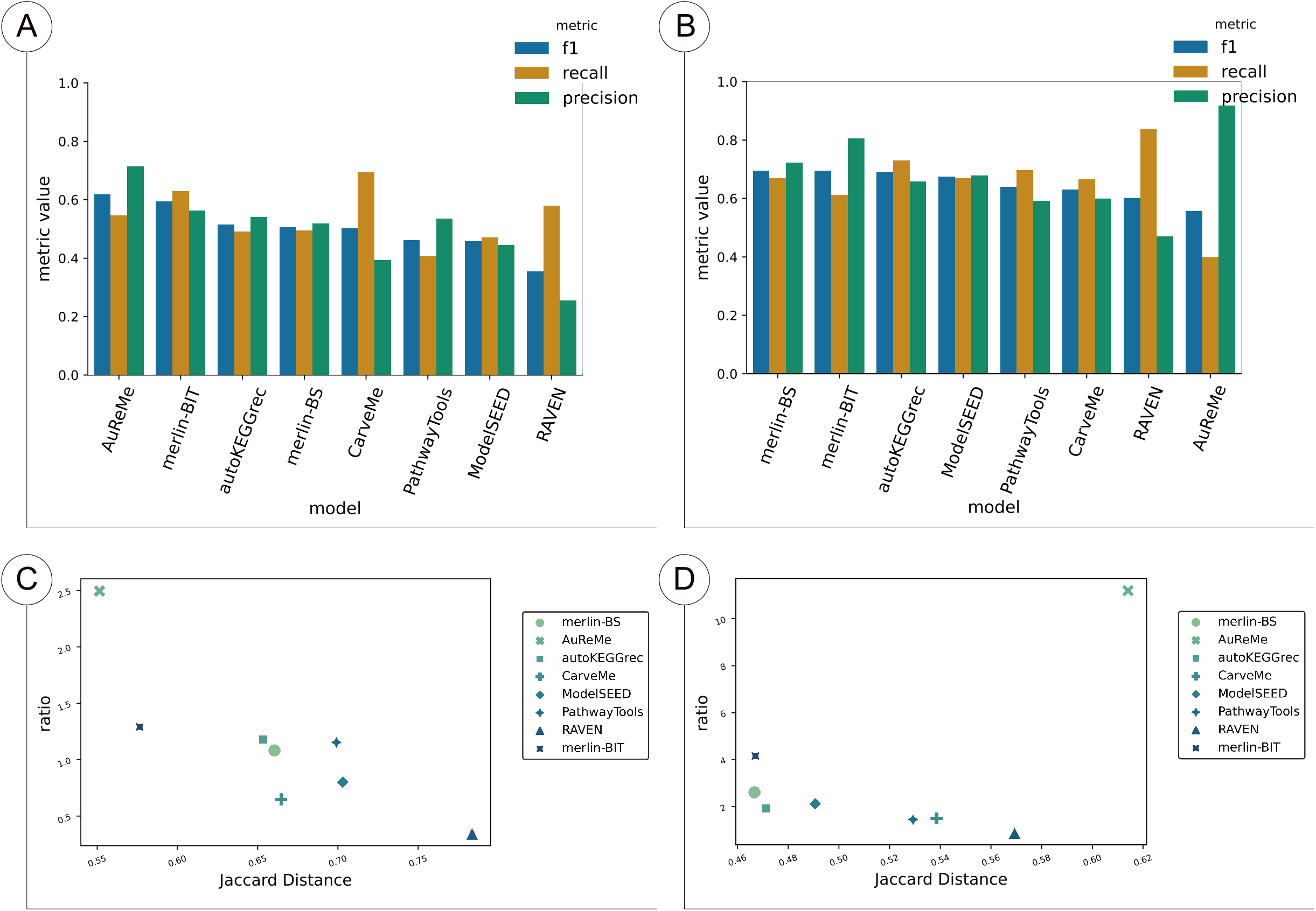
Reaction and gene sets of the draft models of *B. pertussis*. A and C show the results of the reactions, whereas B and D show the results of the genes. A and B enumerate the F1, Recall, and Precision for all tools and methods, sorted by F1 score. C and D depicts the results of TP/FP Ratio as a function of JD.

The reaction F1 scores were considerably low for all models, ranging from 0.33 to 0.49. Figures 3A and 3C indicate that the draft models with better F1, Recall, and JD performances were AuReMe, *merlin-BIT*, and CarveMe. These metrics indicate that the draft models generated by these tools deliver the best balance of TP, FN, and FP. AuReMe achieved the top performance in F1 and JD, whereas CarveMe delivered the higher Recall, although lower F1, JD and Precision than *merlin-BIT*. Also, PathwayTools obtained the best Precision and TP/FP Ratio, although only 379 out of 1299 of the reactions were converted (Table S5 of Supplementary data 2).

Though higher than the reaction sets’ F1 scores, the gene sets’ F1 values were still low for all models, ranging from 0.52 to 0.65. Figure 3B and 3D show that the tools with better F1, Recall, and JD performances were CarveMe, *merlin-BS*, and *merlin-BIT*. RAVEN delivered the best Recall, although it performed poorly for F1 and Precision. On the contrary, AuReMe delivered the best Precision to the detriment of the Recall, which was the worst of all draft models.

##### Lactobacilus plantarum

Regarding the assessment of the models reconstructed for *L. plantarum*, Figure 4 shows all the metrics results for reactions (Figure 4A and 4C), and genes (Figure 4B and 4D). These results are detailed in Tables S7 and S8 (Supplementary data 2).

**Figure 4.**
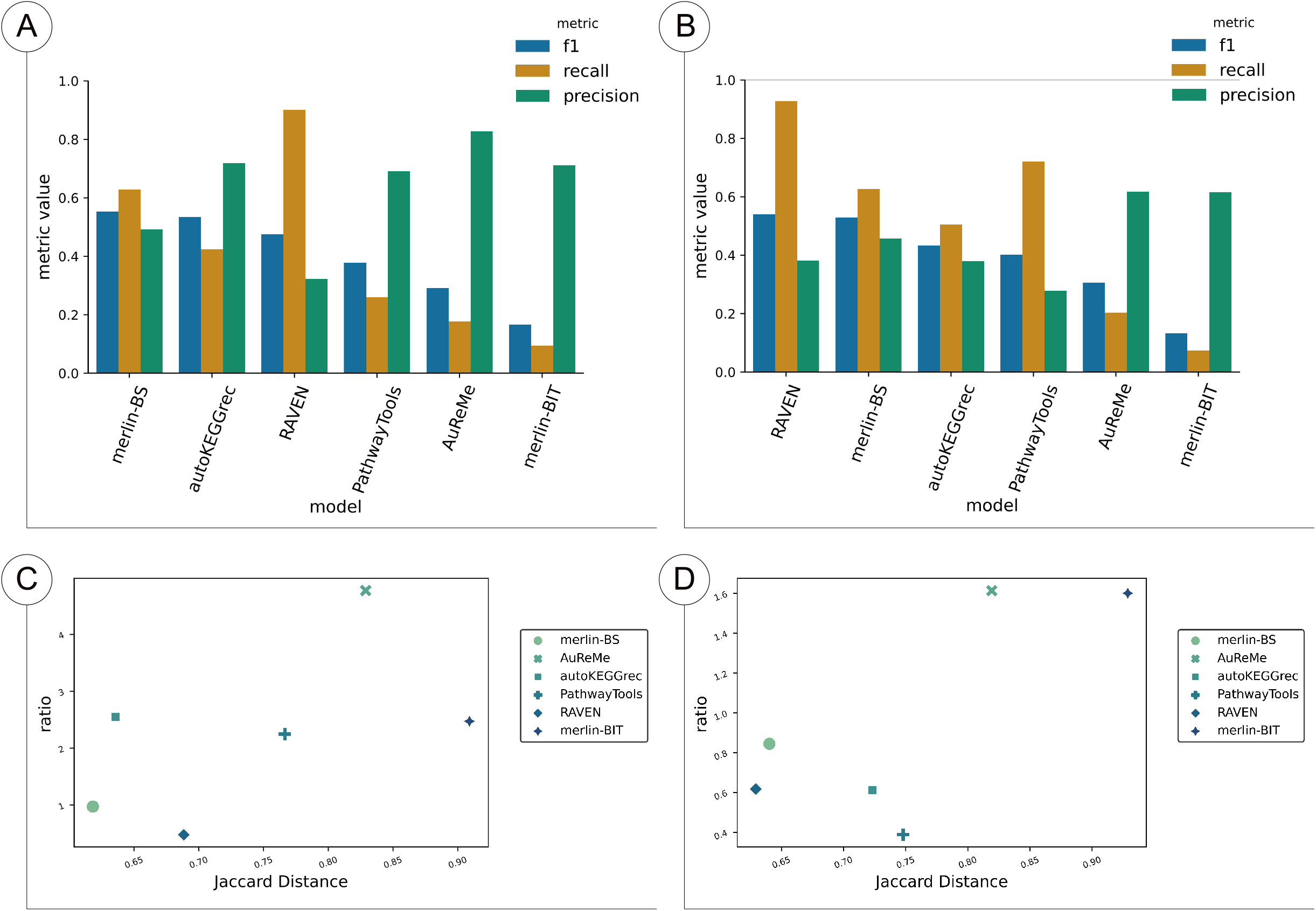
Reaction and gene sets of the draft models of *L. plantarum*. A and C show the results of the reactions, whereas B and D show the results of the genes. A and B enumerate the F1, Recall, and Precision for all tools and methods, sorted by F1 score. C and D depicts the results of TP/FP Ratio as a function of JD.

The F1 score was low and ranged from 0.35 to 0.61 for reaction sets. The AuReMe model of *L. plantarum* obtained the best performance of F1, JD, Precision, Ratio, and Precision regarding reactions. On the other hand, CarveMe delivered the best Recall. AuReMe outperformed all other models in all metrics, followed by *merlin-BIT*, while *merlin-BS* surpassed CarveMe, PathwayTools, ModelSEED, and RAVEN in terms of F1 and JD. Furthermore, it is worth noting that two out of the four top-performing models, regarding F1 and JD, were generated with *merlin*.

Regarding the genes present in the models, *merlin-BS* obtained the best F1 and JD, RAVEN the best Recall, but the worst Precision, and AuReMe the best Precision and TP/FP Ratio. Nonetheless, the F1 values of gene sets were still low, as ranged from 0.55 to 0.69.

##### Toxoplasma gondii

Regarding the assessment of the models reconstructed for *T. gondii*, Figure 5 shows all the metrics for reactions (Figure 5A and 5C), and genes (Figure 5B and 5D). These results are detailed in Tables S9 and S10 (Supplementary data 2).

**Figure 5.**
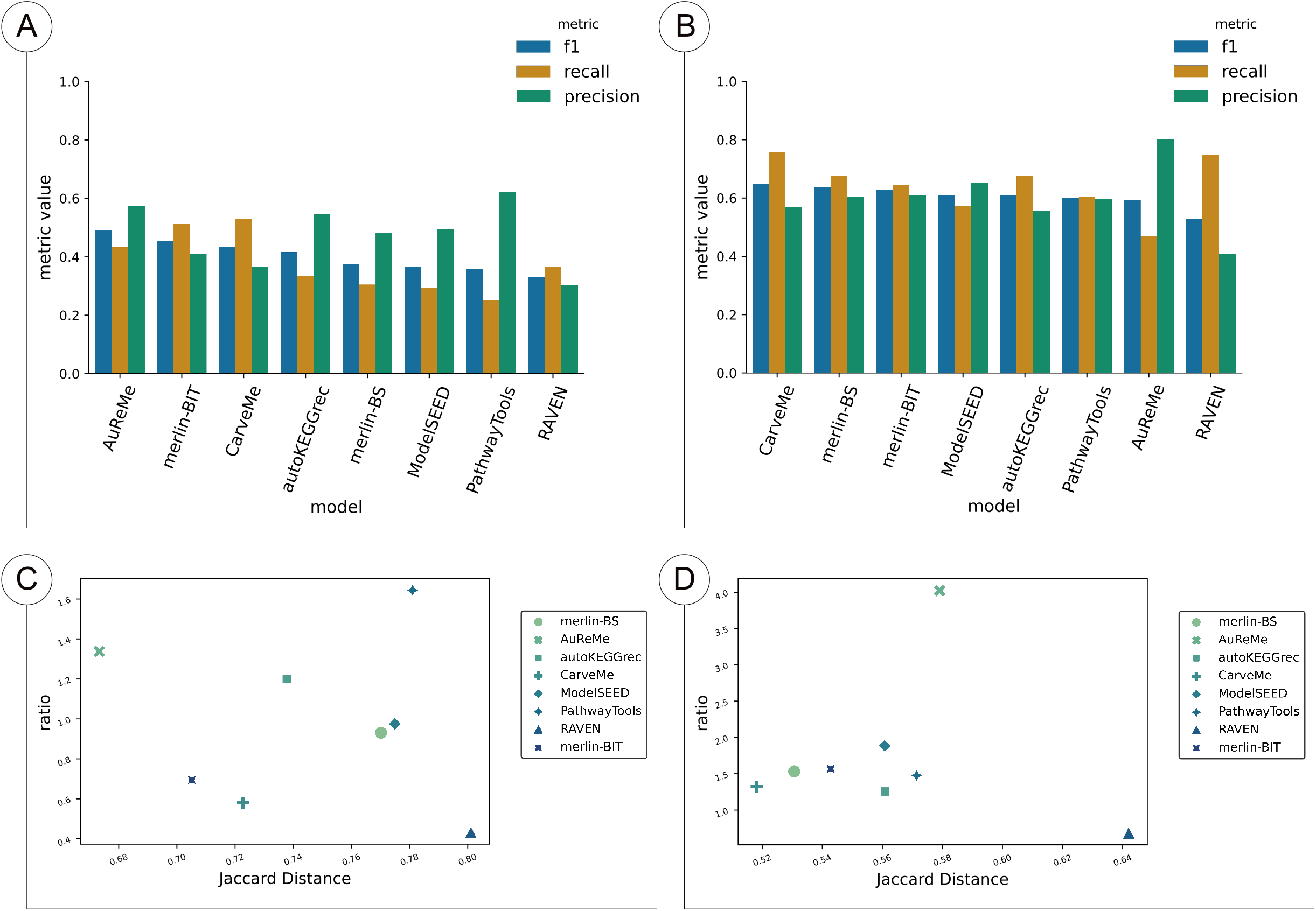
Reaction and gene sets of the draft models of *T. gondii*. A and C show the results of the reactions, whereas B and D show the results of the genes. A and B enumerate the F1, Recall, and Precision for all tools and methods, sorted by F1 score. C and D depicts the results of TP/FP Ratio as a function of JD.

The *T.gondii* results revealed that the three top-performing models in terms of reactions were *merlin-BS*, AutoKEGGRec, and RAVEN, sorted by F1 score. It is worth noting that F1 values ranged from 0.16 to 0.55, which is considerably low. As shown in Table S9 (Supplementary data 2), *merlin-BIT’s* model could only associate 564 reactions, of which 106 were not considered (sink, transport, demand or exchange reactions) and 134 converted to KEGG identifiers. Likewise, AuReMe only included 432 reactions, of which 89 were not considered and 209 converted. AutoKEGGRec also delivered a model with only 567 reactions, of which 564 were present in MetaNetX. Although achieving higher Precision and TP/FP Ratio, these models performed very poorly regarding JD, Recall, and F1.

The F1 and JD of the gene sets revealed that the curated model’s nearest drafts were *merlin-BIT* and merlin-BS. Although RAVEN and *merlin-BS* were the best regarding F1 and JD, as for the reactions results, AuReMe and *merlin-BIT* performed well in Precision and TP/FP Ratio but poorly for the other metrics. These models only included 243 and 104 genes, respectively, obtaining an extremely high number of FN but a low number of FP. F1 values for gene sets ranged from 0.13 to 0.54, which is extremely low.

##### Overall performance

We tested several approaches to generate draft models with *merlin* and validated them with curated and published GSMMs. Overall, reconstructing draft GSMMs of prokaryotes indicated that *merlin-BIT* outperforms the other non-template-based methods, including both *merlin* and the other tools models. Within the non-template-based models generated with *merlin*, the ones generated from BLAST annotations performed marginally better for reaction sets than Diamond’s. That was the case for gene sets concerning *L. plantarum* models, yet not for *B. pertussis*, where Diamond sensitive mode generated the best results. Nonetheless, *SamPler* allows the user to generate slightly better models for almost all cases. Non-template-based models derived from different approaches within *merlin* were not significantly different, delivering metrics that differed very slightly.

As expected, models generated using the biochemical information within the BiGG database were closer to the curated models of *L. plantarum* and *B. pertussis* in terms of included reactions. These results show that the reconstructions depend highly on the information collected from databases. However, this is not true for gene sets, as the models with a better performance were not necessarily those derived from models within BiGG database.

The performance of the reconstruction tools for the *T. gondii* case was substantially different than for prokaryotes. All template-based models (AuReMe and *merlin-BIT*), except RAVEN, obtained poor results and were extremely small compared to the curated model. On the contrary, the non-template-based models performed better. In this sense, the models from *merlin* performed better regarding reactions’ JD and F1 than all others.

Hence, regarding *T. gondii*, template-based models were outperformed. Notably, as *T. gondii* is an eukaryote and an organism substantially less studied than *L. plantarum*, *B. pertussis*, and other prokaryotes, its metabolic network is expectedly more challenging to predict. In this sense, having alternative strategies to address this challenge without depending on the existence/abundance of taxonomically related organisms GSMMs is undoubtedly crucial.

In the context of the state-of-the-art tools, *merlin* improved in terms of TP/FP Ratio and JD for both reactions and genes compared with the previous version evaluated in (4). Such improvements resulted from implementing the new genome and transporters annotation tools.

Overall, the draft models obtained unsatisfactory performances for F1 and JD, the most critical metrics, when considering the whole set of TP, FP, and FN. Such findings corroborate the importance of manually curating a draft GSMM and integrating expert knowledge into metabolic models. *merlin* provides a suitable platform to enhance and facilitate metabolic network curation.

#### Parameters to assess reconstruction tools

We assessed the main features of each of the state-of-the-art tools approached here. Table 1 provides a heatmap of each enumerated feature’s absence, presence or incompletion.

**Table 1.**
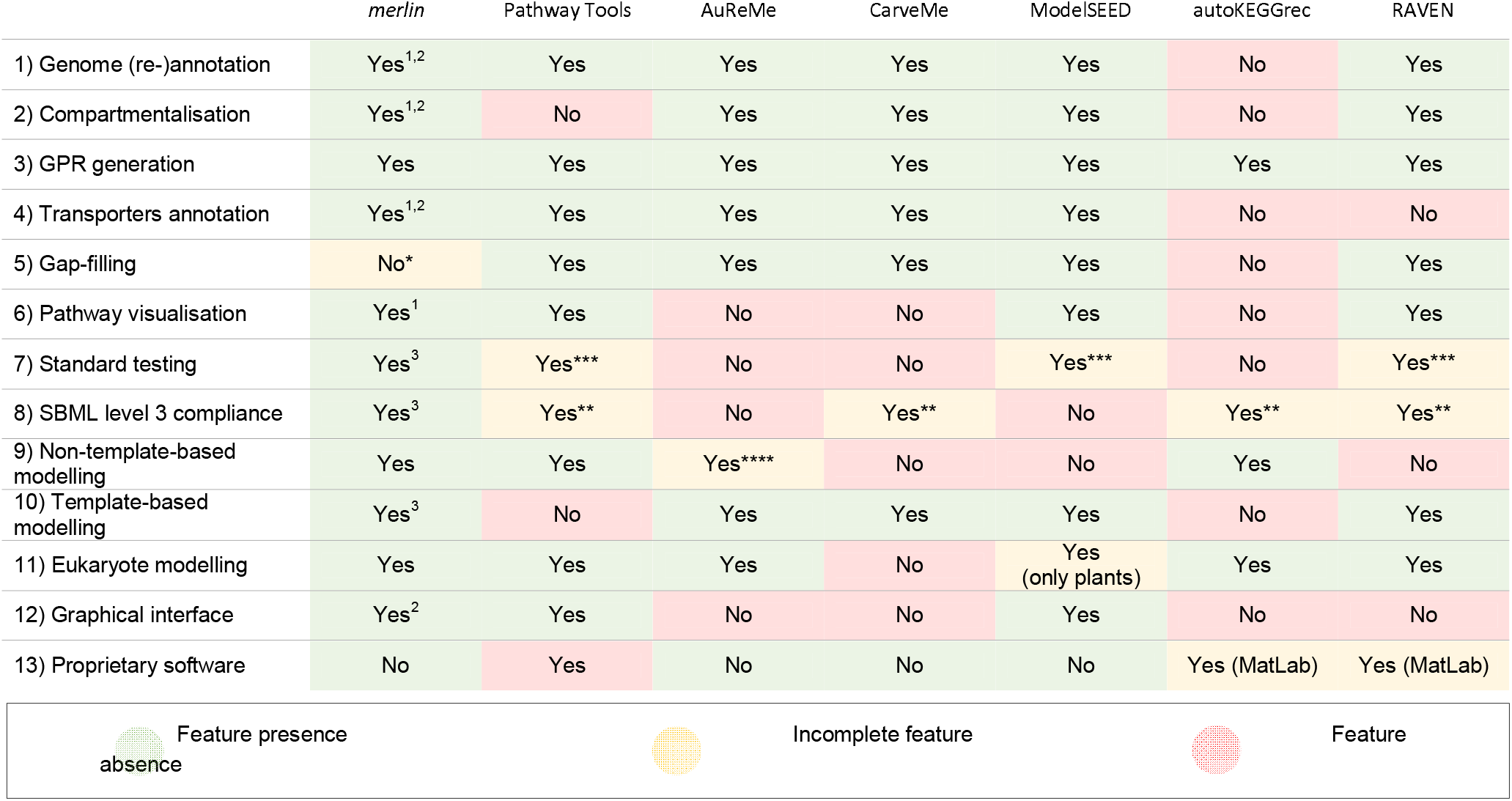
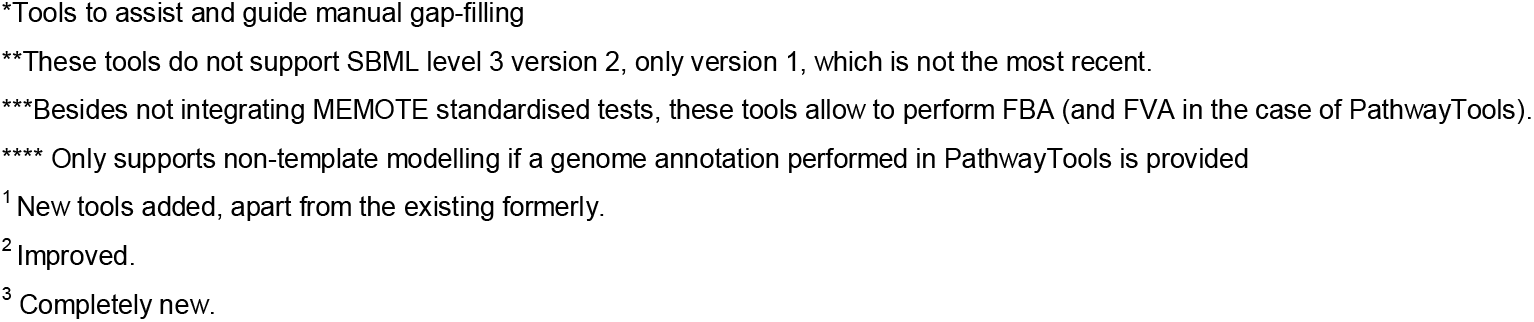
Reconstruction features of each state-of-the-art tool. Green shaded elements indicate the presence/positiveness of the feature, yellow shaded indicate that the feature is incomplete and red shaded ones represent the absence of the feature.

|  | *merlin* | Pathway Tools | AuReMe | CarveMe | ModelSEED | autoKEGGrec | RAVEN |
| --- | --- | --- | --- | --- | --- | --- | --- |
| 1) Genome (re-)annotation | Yes<sup>1,2</sup> | Yes | Yes | Yes | Yes | No | Yes |
| 2) Compartmentalisation | Yes<sup>1,2</sup> | No | Yes | Yes | Yes | No | Yes |
| 3) GPR generation | Yes | Yes | Yes | Yes | Yes | Yes | Yes |
| 4) Transporters annotation | Yes<sup>1,2</sup> | Yes | Yes | Yes | Yes | No | No |
| 5) Gap-filling | No* | Yes | Yes | Yes | Yes | No | Yes |
| 6) Pathway visualisation | Yes<sup>1</sup> | Yes | No | No | Yes | No | Yes |
| 7) Standard testing | Yes<sup>3</sup> | Yes*** | No | No | Yes*** | No | Yes*** |
| 8) SBML level 3 compliance | Yes<sup>3</sup> | Yes** | No | Yes** | No | Yes** | Yes** |
| 9) Non-template-based modelling | Yes | Yes | Yes**** | No | No | Yes | No |
| 10) Template-based modelling | Yes<sup>3</sup> | No | Yes | Yes | Yes | No | Yes |
| 11) Eukaryote modelling | Yes | Yes | Yes | No | Yes<br>(only plants) | Yes | Yes |
| 12) Graphical interface | Yes<sup>2</sup> | Yes | No | No | Yes | No | No |
| 13) Proprietary software | No | Yes | No | No | No | Yes (MatLab) | Yes (MatLab) |

Feature presence
Incomplete feature
Feature
\*Tools to assist and guide manual gap-filling
\*\*These tools do not support SBML level 3 version 2, only version 1, which is not the most recent.
\*\*\*Besides not integrating MEMOTE standardised tests, these tools allow to perform FBA (and FVA in the case of PathwayTools).
\*\*\*\* Only supports non-template modelling if a genome annotation performed in PathwayTools is provided
<sup>1</sup> New tools added, apart from the existing formerly.
<sup>2</sup> Improved.

Table 1 denotes that all the reconstruction tools allow genome annotation except autoKEGGRec, as it imports the genome annotation directly from KEGG. Such dependence can be particularly limiting because the reconstruction cannot be performed when the genome annotation is not present in KEGG. *merlin* has significantly improved and added alternatives for genome annotation and hastened the process of obtaining alignment results for large-sized databases such as UniProtKB, a unique feature among the tools herein enumerated.

Almost all tools can predict or integrate the sub-cellular localisation of enzymes and reactions, except PathwayTools and autoKEGGRec. However, only *merlin* and RAVEN can integrate *de novo* information without propagating the existing annotation from template models. Still, RAVEN only uses *WolfPSORT*, whereas *merlin* provides parsers for WolfPSORT, PSORTb3, and, more recently, LocTree3.

Concerning the transporters annotation and transport reactions integration, the only tools without these features are autoKEGGrec and RAVEN. However, only ModelSEED, *merlin*, and PathwayTools can annotate *de novo* information on transport proteins and complexes and assign reactions to those protein systems. Other tools, namely CarveMe and AuReMe, propagate the transporter annotations from template models. This fact limits the transporters’ annotation to existing GSMMs. On the contrary, *TranSyT* uses an up-to-date version of the manually curated TCDB to annotate transport proteins, whose annotation level is not dependent on the pace of new GSMMs reconstruction.

Pathway visualisation tools can be crucial for curating a metabolic network manually. Only AuReMe, CarveMe, and autoKEGGRec do not support pathway visualisation. PathwayTools and RAVEN provide their approach for presenting the whole metabolic network, while ModelSEED and *merlin* use KEGG and Escher maps. *merlin* delivers direct mappings between the metabolic network within the desktop application and the maps on the web. Moreover, this mapping was also implemented for *MetExploreViz* in the version presented here.

Gap-filling is also a critical step in modelling organisms at the genome-scale, as the current functional annotation of genes is still limited. In this sense, all software tools developed strategies to include approaches to assist and guide manual gap-filling or/and perform it automatically. *merlin* provides tools to find gaps, blocked reactions, and dead-end metabolites using MEMOTE, *BioISO*, or internal algorithms. However, *merlin* cannot perform automatic gap-filling, unlike almost all the other tools, which allow rapidly generating a gapless and simulation-ready model. However, it should be noted that automated gap-filling does not associate reactions with genes, which impairs the quality and eventually the purpose of the model, e.g. hindering the translation of metabolic optimisation strategies to the lab.

Notably, *merlin* is the only tool supporting MEMOTE standard tests and the most recent version of SBML level 3 (version 2) (54). Moreover, *merlin* supports both prokaryote and eukaryote modelling, using model templates or building the metabolic network from scratch.

Finally, *merlin*’s GUI underwent profound changes over the years towards putting user knowledge to the service of genome-scale metabolic modelling, as much as possible. In this sense, *merlin* continues to be an open-access tool and one of the few with a graphical platform to curate the metabolic network.

### *merlin*’s impact on research

Besides the ones published previously, since 2015, tens of high-quality GSMMs have been reconstructed using *merlin*, and at least 11 have been published (55–60). A list of models developed in *merlin* is provided in Table S3 (Supplementary data 1). The applications of these models include synergies with food biotechnology, drug targeting, and efficient biomaterials production. Noticeably, the first genome-wide metabolic model of a ligneous tree was developed using *merlin*. Moreover, *merlin*’s curation tools and interface have been used to (re-)annotate several organisms’ genomes (61–64).

The metabolic annotation of *Pythium irregulare* CBS 494.86’s genome has been employed in *merlin* (61), delivering interesting results towards a better comprehension of Eicosapentaenoic acid production. EC numbers were linked to 3809 genes, and 945 to membrane transporter proteins. Genes associated with amino acid and lipid production, as well as with the consumption of carbon and nitrogen sources present in wastewater, were identified. Such a result provides important insights into the metabolism of *P. irregulare* CBS 494.86 and the possible applications in producing value-added lipids for industrial purposes.

The *Streptococcus pneumoniae* R6 (7) model was reconstructed in *merlin*, with exciting results regarding genome annotation and predictive capability. In this work, 67 essential genes unlisted in the Online GEne Essentiality (OGEE) database (65) were proposed as critical for certain environmental conditions, guiding the discovery of novel drug targets. Moreover, five different studies helped to assess and corroborate the phenotype predictions under different conditions.

Likewise, other pathogens were modelled using *merlin*. A remarkable example was *Candida albicans* (60) whose model now provides an accurate platform for drug targeting. The model correctly predicted 78% of the essential genes (84 out of 108 validated experimentally) under anaerobic growth for different carbon and nitrogen sources.

Four Lactic Acid Bacteria GSMM were recently reconstructed with *merlin;* namely, *Streptococcus thermophilus* LMD-9, *Lactobacillus acidophilus* La-14, *Lactobacillus helveticus* CNRZ32, and *Lactobacillus rhamnosus* GG. The growth rate under different carbon sources, amino acid auxotrophies, and minimal medium were in good agreement with the experimental data. These GSMMs were compared to other models in different environmental conditions for food biotechnology applications (manuscript in preparation).

The *Quercus suber* GSMM (8) epitomises the first genome-scale metabolic model of a ligneous tree. This model comprises 6481 metabolites, 6230 reactions, 7871 genes, and eight different compartments. The authors replicated and simulated growth under autotrophic and heterotrophic conditions, covering the photorespiration process. Furthermore, although not straightforward, this model includes secondary plant metabolism and complete pathways for the bioproduction of suberin monomers, compounds of paramount importance in cork production. Finally, the Quantum Yield and Assimilation Quotient predicted by the model are in accordance with values reported in literature. Another highlight is converting the generic model, reconstructed in *merlin*, into tissue-specific and multitissue models, corroborating the scalability and usability of *merlin* models using Troppo (66).

These studies confirm the reliability and robustness of *merlin*. Moreover, the applications range from food biotechnology to drug targeting and the production of biomaterials, whose relevance is particularly endorsed by several published articles (10–14, 16) and different collaborations. Finally, reconstructing organisms with large genomes, such as *Quercus suber*, highlights *merlin*’s scalability unequivocally.

Since its launch in 2016, *merlin*’s website has been visited more than eleven thousand times, and over three thousand downloads have been performed. *merlin* has gained notoriety over the years and is nowadays recognised as a reference software for reconstructing high-quality GSMM by the scientific community through research articles, reviews, and book chapters, listed in Table S4 of Supplementary data 1.

## CONCLUSION

*merlin* version 4.0 is an updated and robust open-source software developed in Java to reconstruct high-quality GSMMs. Compared with the last published version (*merlin* version 2.0), several improvements and new features were included. These updates are related to the reconstruction process, the software architecture and graphical interface. New semi- and completely automated *plugins* were added, establishing synergies with the improved user-friendly graphical interface to facilitate the model’s curation. Moreover, *merlin*’s software architecture is currently much more modular, allowing the easy insertion of both in-house and third-party computational tools.

Genome functional annotation, draft assembly, model curation, and refinement are critical steps on GSMMs reconstruction and are all covered by *merlin*. The genome functional annotation process with BLAST (22) was considerably accelerated, and a new option is now provided with Diamond (23). Moreover, the alignment results can now be semi- and automatically curated with *SamPler* or the automatic workflow. *TranSyT*, a state-of-the-art tool, performs the transporter systems annotation based on updated versions of TCDB. Notably, template and non-template-based modelling are now supported using BiGG and KEGG information. Finally, the model curation is leveraged by multiple tools for tracking network errors and inconsistencies, complemented by network visualisation add-ons.

The scientific community has extensively used *merlin*, as corroborated by the multiple published works and considerable user-base expansion. The scalability and reliability of *merlin* 4.0 have been showcased with the reconstruction of multiple models, particularly with the *Quercus suber*’s, the first GSMM of a ligneous tree. This study confirmed that manual curation is essential to obtaining high-quality models, as most draft GSMMs are still far from being similar to validated and published ones. In this sense, the enhanced GUI represents an advantage over similar tools with less refined approaches for manual curation. In addition, we showed that *merlin* could generate satisfactory draft models compared with other tools.

In conclusion, *merlin* version 4.0 is an integrated, open-source, and updated platform designed to anchor GSMM’s reconstruction. The trade-off between automatic processes and assisted manual curation ensured by *merlin* allows users to leverage their expertise towards high-quality reconstructions.

## DATA AVAILABILITY

*merlin* is an open-source application currently available for Linux, Windows, and macOS. It is distributed under the GNU General Public License at the website https://www.merlin-sysbio.org. Moreover, *merlin* source code, including plugins, is fully available at https://github.com/merlin4-sysbio. The draft models assessment scripts and files are available at https://github.com/BioSystemsUM/merlinv4_paper.

Comprehensive documentation is provided at https://merlin-sysbio.org/documentation/. Animated snapshots of *merlin’s* interface and clear descriptions of each step in the reconstruction are therein provided.

## SUPPLEMENTARY DATA

Supplementary Data are available at NAR Online.

