## Supplementary data 1 for "*merlin* v4.0: an updated platform for the reconstruction of high-quality genome-scale metabolic models"

* To whom correspondence should be addressed.

### Supplementary data 1 – *merlin's* plugins implementation, databases and models

#### ORM

The new features available in the newest version require the storage and manipulation of complex biological data. In the previous version, the interaction between the MySQL (https://www.mysql.com/) database and the biological data involved prepared statements. Database maintenance and management using prepared statements can be a time-consuming task that requires technical skills and may hinder the development of new features.

To address this problem, two frameworks, Hibernate (https://hibernate.org/) and Spring Data (https://spring.io/projects/spring-data) were integrated into *merlin*'s engine. Hibernate is an open-source ORM framework created to simplify development, allowing the conversion of Plain Old Java Objects (POJO) classes into database tables by using mapping configurations. The data present in a Java object is automatically converted into Structured Query Language (SQL) data without the development of SQL queries, required for the prepared statements. The adoption of an ORM framework improved the database maintenance, as any modification of Java entities can be automatically reflected into the database schema, facilitating the development of database migration patches between versions of *merlin* and its plugins. Another improvement prompted by the ORM adoption was the option to use the H2 database (http://h2database.com/), a Java embedded database, which overrides the technical requirement of installing MySQL in the machine that will run *merlin*.

Spring data was integrated into *merlin's* engine to foster Hibernate framework functionalities and further improve *merlin*'s database management. In the new version of *merlin*, Spring-based data accesses manage the database operations performed by Hibernate ORM and *merlin* CORE data structures. The data accesses split the database operations from *merlin*'s operations, enabling the development of improved data storage schemas, transparent to *merlin*'s operations, which allows integrating different types of database, e.g. non-relational databases like MongoDB (https://www.mongodb.com/) or Neo4J (https://neo4j.com/).

**Table S1 –** Databases used by *merlin.*

| Database / Online resource | Description | Operations using the database | Access | Reference |
|---|---|---|---|---|
| Kyoto Encyclopedia of Genes and Genomes (KEGG) | KEGG is an integrated resource encompassing 16 different databases divided by categories. These categories are divided into Systems, Genomic, Chemical, and Health information. The Systems category comprises pathways, modules, as well as relevant hierarchies of different biological entities. The Genomic category includes genes, genomes, as well as valuable genome annotations. The Chemical one compiles metabolic and biochemical information relevant for building up a metabolic network. The Health category includes drugs and health related substances. | • Importing metabolic data;<br>• Performing GPR associations;<br>• "Draw in Browser" operation. | Rest API through http://rest.kegg.jp/ | (1) |
| National Center for Biotechnology Information (NCBI) | NCBI comprises databases with vast application in biomedicine and biotechnology, including bioinformatics tools with free usage. It houses a database for multiple genomes from all domains of life, as well as a taxonomic hierarchy regularly updated, providing relevant information in the context of metabolic modelling. | • Importing genomes<br>• Retrieving taxonomy lineage | Using Entrez taxonomy and Entrez Proteins databases through https://eutils.ncbi.nlm.nih.gov/entrez/eutils | (2) |
| ModelSEED | ModelSEED is an integrated database that includes information from different databases and GSM models. The biochemical data therein present involves reactions' and metabolites' compartmentalization, transport reactions, charged molecules and proton balancing. Moreover, annotations and database's cross-references are provided. | • Importing metabolic data<br>• Biomass Templates<br>• Cross-references for metabolites and reactions | Files download from https://github.com/ModelSEED/ModelSEEDDatabase | (3) |
| Transporter Classification Database (TCDB) | TCDB is a freely available resource that accounts for the enumeration and curated characterization of transporter systems from varied types of living organisms. It includes a classification system for transporters that can incorporate functional and phylogenetic information. This system is approved and adopted by the International Union of Biochemistry and Molecular Biology (IUBMB). | • Using TranSyT for transporter systems annotation | Using TranSyT's auto-updatable TCDB's database | (4) |
| Universal Protein Resource Knowledgebase (UniProtKB) | UniProtKB is a knowledgebase of functional information on proteins. It includes two main databases: Swiss-Prot and TrEMBL. The former corresponds to manually curated information, whereas the later contains unreviewed and automatically generated data. | • Reference database to perform the enzymes annotation | Through EBI's remote access | (5) |

| | | | | |
|---|---|---|---|---|
| Gastric Cancer Platform | GCPlatform is a novel knowledge-base and web portal to foster scientific discovery and drug development efforts in gastric cancer research, developing novel methods to integrate data from different origins, results of data analysis, and data mining tools. It houses protein information from multiple sources (e.g. Uniprot, ProteomicsDB, InterPro) and provides summarized protein information with repository cross-referencing. | • Identifier mapping and enzyme information retrieval after performing alignments | Rest API through https://gcplatform.bio.di.uminho.pt/api/prot/find/ids | - |

**Table S2 –** Merlin's plugins' implementation and references.

| Reconstruction step | Tool | Implementation | Third-party software | Reference |
|---|---|---|---|---|
| Genome annotation | *SamPler** | Integrated in *merlin* | no | (6) |
| | Automatic workflow* | Integrated in *merlin* | no | - |
| | *TranSyT** | Remote access to *TranSyT*'s RestAPI and parsers to integrate the results; | no | (7) |
| | Comparmentalization* | Web-scrapers for *LocTree3* (https://rostlab.org/services/loctree3/) and *WolfPSort* (https://wolfpsort.hgc.jp/) reports' webpage, and report parser for *PSort3b* (https://www.psort.org/psortb/) | yes | (8-10) |
| Model reconstruction | GPR associations | Integrated in *merlin*, using KEGG API | no | (11) |
| | Biomass formulation* | Integrated in *merlin* | no | (12) |
| | Draft Reconstruction* | Integrated in *merlin* | no | - |
| | Draft Reconstruction* | BiGG Integration Tool - Access to an internal RestAPI – running in BioSystems servers | no | (13) |
| Curation | *BioISO** | Remote access to *BioISO*'s RestAPI and parsers to render the results | no | (14) |
| | *MEMOTE** | Access to an internal RestAPI – running in BioSystems servers – and parser to render the results | yes | (15) |
| | *EscherMaps** | Access to an internal RestAPI – running in BioSystems servers – opening results HTML in user's default browser | yes | (16) |
| | *MetExploreViz** | Web-service run locally - remote access to *MetExploreViz*'s RestAPI | yes | (17) |

* New features of version 4.0

**Table S3 -** Published models reconstructed in merlin.

| Type | Organism | Applications | Metabolites | Reactions | Genes | Compartments | Remarks | Reference |
|---|---|---|---|---|---|---|---|---|
| Gram-negative bacteria | *Helicobacter pylori 26695* | Pathogen | 412 | 640 | 383 | 3 | 191 new metabolic functions were identified in the genome re-annotation. Better phenotypical predictions were obtained in the specific growth rate comparing to other models reconstructed with other tools. Agreement with experiments using glutamate as sole carbon source, whereas its analogous were not capable of predicting growth under glutamate consumption. As for the gene essentiality, it performed slightly better than its analogous, using experimental data to corroborate it. | (18) |
| | *Streptococcus pneumoniae R6* | Pathogen | 489 | 596 | 372 | 2 | First model for any *S. pneumoniae* strain. Complete functional annotation of the genome: 372 metabolic genes identified out of the 2046 candidates. Accurate phenotypical predictions under different environmental conditions, different carbon sources and oxygen availability, being corroborated by 5 different studies. 67 essential genes not listed in OGEE database were identified. | (19) |
| | *Actinobacillus succinogenes 130Z* | Bioproduction of succinic acid | 713 | 1072 | 722 | - | Provided a better comprehension of *A. succinogenes* metabolism and capacities. Naturally, this organism is one of the major producers of succinic acid and the reconstructed model presents advantages over other SA-producing strains, such as its ability to consume low-cost carbon sources and C5, C6 sugars. | (20) |
| Gram-positive bacteria | *Streptococcus thermophilus LMD-9* | Food biotechnology | 794 | 962 | 431 | 2 | Allowed the comparison of their fermentative behaviour in different environmental conditions with other Lactic Acid Bacteria. | (21) |
| | *Lactobacillus acidophilus La-14* | Food biotechnology | 574 | 677 | 470 | 2 | | |
| | *Lactobacillus helveticus CNRZ32* | Food biotechnology | 562 | 618 | 389 | 2 | | |
| | *Lactobacillus rhamnosus GG* | Food biotechnology | 740 | 876 | 651 | 2 | | |
| | *Enterococcus faecalis V583* | Pathogen | 576 | 603 | 366 | 2 | First model of this species. Phenotypical predictions were in accordance with results from chemostat experiments, being able to accurately predict all the organism's end-products comparing to the distribution of fluxes observed in vivo. | (22) |

Continued on the next page.

**Table S3 –** Continued from the previous page.

| Type | Organism | Applications | Metabolites | Reactions | Genes | Compartments | Remarks | Reference |
|---|---|---|---|---|---|---|---|---|
| Fungi | *Kluyveromyces lactis* | Food biotechnology | 1476 | 1867 | 906 | 4 | First model of this species: 45 genes updated from the previous genome annotation, 22 new metabolic genes were identified. In silico growth predictions in agreement with Biolog experiments and the online catalogue of strains CBS-KNAW. Accurate predictions using 9 different carbon sources. Replication of 90% of the knockouts from several experiments published over the last three decades. | (23) |
| | *Candida albicans* | Pathogen | 926 | 1221 | 781 | 4 | First model of this species. It correctly predicted 78% of the identified essential genes (84 out of 108 validated experimentally). Proved to be accurate prediction of anaerobic growth utilizing different carbon and nitrogen sources. | (24) |
| Plantae | *Quercus suber* | Cork production | 6481 | 6230 | 7871 | 8 | First model of a ligneous tree. Besides the generic model, were also obtained tissue-specific models, for leaf, inner bark, and phellogen, and a diel multi-tissue model. | (25) |

Table S4 – *merlin's* research impact as of 2015.

| Authors | year | Title | Journal/Book/Conference |
|---|---|---|---|
| **Machado,D., Herrgård,M.J. and Rocha,I.** | 2016 | Stoichiometric Representation of Gene–Protein–Reaction Associations Leverages Constraint-Based Analysis from Reaction to Gene-Level Phenotype Prediction | PLOS Computational Biology |
| **Jouhten,P., Boruta,T., Andrejev,S., Pereira,F., Rocha,I. and Patil,K.R.** | 2016 | Yeast metabolic chassis designs for diverse biotechnological products | Scientific Reports |
| **Pfau,T., Christian,N., Masakapalli,S.K., Sweetlove,L.J., Poolman,M.G. and Ebenhöh,O.** | 2016 | The intertwined metabolism of Medicago truncatula and its nitrogen fixing symbiont Sinorhizobium meliloti elucidated by genome-scale metabolic models | bioRxiv |
| **Weber,T. and Kim,H.U.** | 2016 | The secondary metabolite bioinformatics portal: Computational tools to facilitate synthetic biology of secondary metabolite production | Synthetic and Systems Biotechnology |
| **Kim,W.J., Kim,H.U. and Lee,S.Y.** | 2017 | Current state and applications of microbial genome-scale metabolic models | Current Opinion in Systems Biology |
| **Hädicke,O. and Klamt,S.** | 2017 | EColiCore2: a reference network model of the central metabolism of Escherichia coli and relationships to its genome-scale parent model | Scientific Reports |
| **Lopes,H. and Rocha,I.** | 2017 | Genome-scale modeling of yeast: chronology, applications and critical perspectives | FEMS Yeast Research |
| **102. Thor,S., Peterson,J.R. and Luthey-Schulten,Z.** | 2017 | Genome-Scale Metabolic Modeling of Archaea Lends Insight into Diversity of Metabolic Function | Archaea |
| **Chiappino-Pepe,A., Pandey,V., Ataman,M. and Hatzimanikatis,V.** | 2017 | Integration of metabolic, regulatory and signaling networks towards analysis of perturbation and dynamic responses | Current Opinion in Systems Biology |
| **Mendes-Soares,H. and Chia,N.** | 2017 | Community metabolic modeling approaches to understanding the gut microbiome: Bridging biochemistry and ecology | Free Radical Biology and Medicine |
| **Fondi,M. and Fani,R.** | 2017 | Constraint-based metabolic modelling of marine microbes and communities | Marine Genomics |
| **Karlsen,E., Schulz,C. and Almaas,E.** | 2018 | Automated generation of genome-scale metabolic draft reconstructions based on KEGG | BMC Bioinformatics |
| **Lakshmanan,M., Koduru,L. and Lee,D.-Y.** | 2018 | Software Applications for Phenotype Analysis and Strain Design of Cellular Systems | In Emerging Areas in Bioengineering |

| | | | |
|---|---|---|---|
| **100. Muller,E.E.L., Faust,K., Widder,S., Herold,M., Martínez Arbas,S. and Wilmes,P.** | 2018 | Using metabolic networks to resolve ecological properties of microbiomes | Current Opinion in Systems Biology |
| **Rau,M.H. and Zeidan,A.A.** | 2018 | Constraint-based modeling in microbial food biotechnology | Biochemical Society Transactions |
| **Machado,D., Andrejev,S., Tramontano,M. and Patil,K.R.** | 2018 | Fast automated reconstruction of genome-scale metabolic models for microbial species and communities | Nucleic Acids Research |
| **Choi,K.R., Jang,W.D., Yang,D., Cho,J.S., Park,D. and Lee,S.Y.** | 2019 | Systems Metabolic Engineering Strategies: Integrating Systems and Synthetic Biology with Metabolic Engineering | Trends in Biotechnology |
| **101. Angione,C.** | 2019 | Human Systems Biology and Metabolic Modelling: A Review—From Disease Metabolism to Precision Medicine | BioMed Research International |
| **Castillo,S., Patil,K.R. and Jouhten,P.** | 2019 | Yeast Genome-Scale Metabolic Models for Simulating Genotype–Phenotype Relations | Yeasts in Biotechnology and Human Health |
| **Gu,C., Kim,G.B., Kim,W.J., Kim,H.U. and Lee,S.Y.** | 2019 | Current status and applications of genome-scale metabolic models | Genome Biology 2019 20:1 |
| **Santibáñez,R., Garrido,D. and Martin,A.J.M.** | 2020 | Atlas : automatic modeling of regulation of bacterial gene expression and metabolism using rule-based languages | Bioinformatics |
| **Alballa,M., Aplop,F. and Butler,G.** | 2020 | TranCEP: Predicting the substrate class of transmembrane transport proteins using compositional, evolutionary, and positional information | PLOS ONE |
| **Lam,T.J., Stamboulian,M., Han,W. and Ye,Y.** | 2020 | Model-based and phylogenetically adjusted quantification of metabolic interaction between microbial species | PLOS Computational Biology |
| **Simensen,V., Voigt,A. and Almaas,E.** | 2020 | High-quality genome-scale metabolic model of Aurantiochytrium sp. T66 | bioRxiv |
| **Little,B.J., Blackwood,D.J., Hinks,J., Lauro,F.M., Marsili,E., Okamoto,A., Rice,S.A., Wade,S.A. and Flemming,H.-C.** | 2020 | Microbially influenced corrosion—Any progress? | Corrosion Science |
| **Xu,X., Liu,Y., Du,G. and Liu,L.** | 2020 | Systems biology, synthetic biology, and metabolic engineering | In Systems and Synthetic Metabolic Engineering |
| **Seif,Y. and Palsson,B.Ø.** | 2021 | Path to improving the life cycle and quality of genome-scale models of metabolism | Cell Systems |

| | | | |
|---|---|---|---|
| **Esvap,E. and Ulgen,K.O.** | 2021 | Advances in Genome-Scale Metabolic Modeling toward Microbial Community Analysis of the Human Microbiome | ACS Synthetic Biology |
| **Kuriya,Y., Inoue,M., Yamamoto,M., Murata,M. and Araki,M.** | 2021 | Knowledge extraction from literature and enzyme sequences complements FBA analysis in metabolic engineering | Biotechnology Journal |
| **Moscardó García,M., Pacheco,M., Bintener,T., Presta,L. and Sauter,T.** | 2021 | Importance of the biomass formulation for cancer metabolic modeling and drug prediction | iScience |
| **Saraiva,J.P., Worrich,A., Karakoç,C., Kallies,R., Chatzinotas,A., Centler,F. and Nunes da Rocha,U.** | 2021 | Mining Synergistic Microbial Interactions: A Roadmap on How to Integrate Multi-Omics Data | Microorganisms |
| **Chung,W.Y., Zhu,Y., Mahamad Maifiah,M.H., Shivashekaregowda,N.K.H., Wong,E.H. and Abdul Rahim,N.** | 2021 | Novel antimicrobial development using genome-scale metabolic model of Gram-negative pathogens: a review | The Journal of Antibiotics |
| **Patra,P., Das,M., Kundu,P. and Ghosh,A.** | 2021 | Recent advances in systems and synthetic biology approaches for developing novel cell-factories in non-conventional yeasts | Biotechnology Advances |
| **Sangha,G.S., Goergen,C.J., Prior,S.J., Ranadive,S.M. and Clyne,A.M.** | 2021 | Preclinical techniques to investigate exercise training in vascular pathophysiology | American Journal of Physiology-Heart and Circulatory Physiology |
| **Frades,I., Foguet,C., Cascante,M. and Araúzo-Bravo,M.J.** | 2021 | Genome Scale Modeling to Study the Metabolic Competition between Cells in the Tumor Microenvironment | Cancers |
| **Wendering,P. and Nikoloski,Z.** | 2022 | COMMIT: Consideration of metabolite leakage and community composition improves microbial community reconstructions | PLOS Computational Biology |
| **Camborda,S., Weder,J.-N. and Töpfer,N.** | 2022 | CobraMod: a pathway-centric curation tool for constraint-based metabolic models | Bioinformatics |
| **Chu,L., Li,S., Dong,Z., Zhang,Y., Jin,P., Ye,L., Wang,X. and Xiang,W.** | 2022 | Mining and engineering exporters for titer improvement of macrolide biopesticides in Streptomyces | Microbial Biotechnology |

| | | | |
|---|---|---|---|
| **Carey,M.A., Medlock,G.L., Stolarczyk,M., Petri,W.A., Guler,J.L. and Papin,J.A.** | 2022 | Comparative analyses of parasites with a comprehensive database of genome-scale metabolic models | PLOS Computational Biology |
| **Pathania,R., Srivastava,A., Srivastava,S. and Shukla,P.** | 2022 | Metabolic systems biology and multi-omics of cyanobacteria: Perspectives and future directions | Bioresource Technology |
| **Saa,P., Urrutia,A., Silva-Andrade,C., Martín,A.J. and Garrido,D.** | 2022 | Modeling approaches for probing cross-feeding interactions in the human gut microbiome | Computational and Structural Biotechnology Journal |
| **Navid,A.** | 2022 | Curating COBRA Models of Microbial Metabolism | Microbial Systems Biology |
