## Supplementary data 2 for "*merlin* v4.0: an updated platform for the reconstruction of high-quality genome-scale metabolic models"

* To whom correspondence should be addressed.

### Supplementary data 2 – draft models' assessment

**Drafts Assessment approach**

These draft reconstructions were compared with manually curated and validated models—the GSMMs of *L. plantarum, B. pertussis,* and *T. gondii* were iLP728, iBP1870, and ToxoNet1, respectively.

iBP1870 contains 770 genes and 1672 reactions. Regarding the latter, 564 (33.73%) are sink, demand, exchange or transport reactions. Out of the remaining 1108, only 933 were converted to other database identifiers.

iLP728 contains 771 reactions and 728 genes. Regarding the latter, 228 (29,57%) were either sink, demand, exchange or transport reactions. Out of the remaining 543, 479 were converted to other database identifiers.

ToxoNet1 is composed of 4414 reactions and 940 genes. Regarding the latter, 3316 (75,12%) were either sink, demand, exchange or transport reactions. Out of the remaining 1098, 957 were converted to other database identifiers.

Table S1 – Models' features

| | Total reactions | Sinks | Exchange | Transport | Demand | Compartments | Genes |
|---|---|---|---|---|---|---|---|
| iBP1870 | 1672 | 0 | 204 | 355 | 5 | 3 | 770 |
| iLP728 | 771 | 0 | 113 | 216 | 1 | 2 | 728 |
| ToxoNet1 | 4414 | 0 | 2056 | 1251 | 0 | 4 | 940 |

### Evaluated Tools

#### AuReMe

AuReMe (Automatic Reconstruction of Metabolic Models) is a Docker-based software that allows reconstructing simulation-ready GSMMs using others as templates or inheriting a genome annotation from Pathway Tools.

The AuReME Docker image version 2.4 was utilised to generate the draft models. AuReMe requires other models as templates and the genome files of the organisms under study, including the template ones. In this sense, the genome FASTA files were preprocessed to contain only the locus tag in the header of each gene. Moreover, the template models were imported to *merlin* in SBML level 3 and exported in SBML level 2 format. Otherwise, AuReMe could not be run as it does not support SBML level 3 template models. Then, an orthology-based reconstruction was generated using the Ortho-Finder algorithm.

#### PathwayTools

Pathway Tools is software designed for the development of organism-specific metabolic databases. The application is developed in Lisp and provides a graphical interface that allows the user to visualise the whole constructed network. Moreover, it includes the assisted analysis of experimental data in the metabolic network.

The Pathway Tools version 25.5 was downloaded and utilised to generate draft models of the mentioned organisms under study. For this matter, we uploaded the genome file in GenBank format and performed the metabolic network reconstruction with the Pathologic algorithm using default parameters. Moreover, transport reactions were generated and included. Finally, the draft model was exported in an SBML format.

#### CarveMe

CarveMe is a python package focused on fast reconstructing microbial metabolic models. It provides a top-down approach that creates simulation-ready models based on BiGG universal database. This tool uses Diamond to prioritise reactions with stronger genetic evidence (by similarity).

The protein sequences were used as input in FASTA format using default parameters, providing simulation-ready GSM models in SBML format. The gap-filling mode of this tool was not used.

**RAVEN**

RAVEN (Reconstruction, Analysis and Visualization of Metabolic Networks) is a command-line tool compliant with COBRA Toolbox v3, thus running in MATLAB. It provides functionalities to reconstruct models from scratch or use template models, merge networks from different databases and curate from the command line.

The protein FASTA files were used as input to version 2.5.3 of the tool, using the KEGG's "prok90_kegg94" and "euk90_kegg94" templates for prokaryotic and eukaryotic organisms, respectively. The tool was used with default parameters, except for the "keepUndefinedStoich" and "keepIncomplete" (were set as False), and the cutOff value (adjusted to $10^{-30}$).

**ModelSEED**

ModelSEED is a web tool to analyse and reconstruct GSMMs. The genome annotation is performed by RAST within ModelSEED, reconstructing the metabolic network based on that annotation. Moreover, it provides a graphical interface to analyse and gap-fill the model and run simulations.

ModelSEED version 2 (November 2021) was used to reconstruct the draft models of the organisms under study. The genome FASTA files were inputted directly through the web interface, and the template models were defined by each type of organism (gram-positive bacteria for *L. plantarum* and gram-negative bacteria for *B. pertussis*). Finally, we selected a complete medium for the reconstruction.

**AutoKEGGRec**

AutoKEGGRec v1.01 is a command-line tool that runs in MATLAB and provides compliance with COBRA Toolbox v3. This tool is highly suitable to generate multiple GSMMs only in one run and can also generate metabolic models for communities. The reconstruction is dependent on the availability of data for the target organism in KEGG.

The KEGG's organism identifiers for *L. plantarum*, *B. pertussis*, and *T. gondii* were used as input, using the "SingleRecs" mode, suitable for reconstructing GSM models of individual organisms. After running the tool, the models were exported in SBML format.

***merlin***

Regarding *merlin,* we generated several more models to validate all the new and updated features. In this sense, the first step encompassed uploading the genome FASTA file, which was aligned against the Swiss-Prot and TrEBML databases, sequentially, using BLAST or Diamond. The default BLAST and Diamond parameters were inputted; however, two Diamond sensitivity modes were assessed: "fast" and "sensitive". Afterwards, the automatic annotation and *SamPler* were used to annotate the genomes based on the BLAST/Diamond results. The species and genus selected for *automatic annotation* are shown in Tables S1, S2, and S3. The selection of organisms was based on phylogenetic trees obtained by applying a neighbour-joining method to a multiple alignments generated by Clustal Omega of ribosomal RNA (16S for prokaryotes and 18S for eukaryotes) of species well-represented at

Swiss-Prot. The results of this selection are described in Supplementary material 3. Furthermore, transport reactions were generated with *TranSyT*. Finally, GPR rules were constructed using blast+ version 2.11.0.

Table S2 – Automatic workflow's ranking for *B. pertussis*.

| genus/species | organism | e-value threshold | reviewed |
|---|---|---|---|
| species | *Bordetella pertussis* | 1.0E-30 | true |
| genus | *Bordetella* | 1.0E-30 | true |
| species | *Pseudomonas putida* | 1.0E-30 | true |
| species | *Escherichia coli* | 1.0E-30 | true |
| species | *Klebsiella pneumoniae* | 1.0E-30 | true |
| species | any | 1.0E-30 | true |
| species | *Bordetella pertussis* | 1.0E-30 | false |
| genus | *Bordetella* | 1.0E-30 | false |
| species | *Pseudomonas putida* | 1.0E-30 | false |

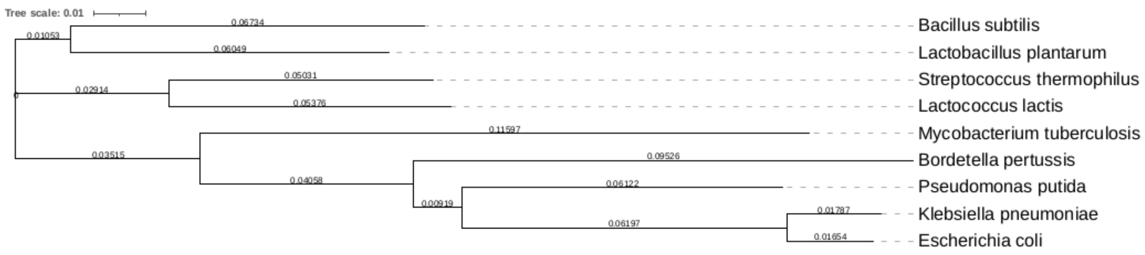


Figure S1 – Phylogenetic tree obtained by applying a neighbour-joining method to a multiple alignment generated by Clustal Omega of 16S ribosomal RNA.

Table S3 – Automatic workflow's ranking for *L. plantarum*.

| genus/species | organism | e-value threshold | reviewed |
|---|---|---|---|
| species | *Lactobacillus plantarum* (strain ATCC BAA-793 / NCIMB 8826 / WCFS1) | 1.0E-30 | true |
| genus | *Lactobacillus* | 1.0E-30 | true |
| species | *Bacillus subtilis* | 1.0E-30 | true |
| species | *Streptococcus thermophilus* | 1.0E-30 | true |
| species | *Lactococcus lactis* | 1.0E-30 | true |
| species | any | 1.0E-30 | true |
| species | *Lactobacillus plantarum (strain ATCC BAA-793 / NCIMB 8826 / WCFS1)* | 1.0E-30 | false |
| genus | *Lactobacillus* | 1.0E-30 | false |
| species | *Bacillus subtilis* | 1.0E-30 | false |

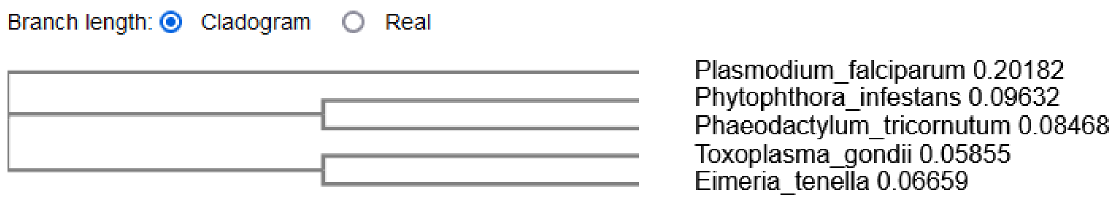


Figure S2 – Phylogenetic tree obtained by applying a neighbour-joining method to a multiple alignment generated by Clustal Omega of 18S ribosomal RNA.

Table S4 – Automatic workflow's ranking for *T. gondii*.

| **genus/species** | **organism** | **e-value threshold** | **reviewed** |
|---|---|---|---|
| species | *Toxoplasma gondii* ME49 | 1.0E-30 | true |
| genus | *Toxoplasma* | 1.0E-30 | true |
| species | *Eimeria tenella* | 1.0E-30 | true |
| species | *Phaeodactylum tricornutum* | 1.0E-30 | true |
| species | *Plasmodium falciparum* | 1.0E-30 | true |
| species | any | 1.0E-30 | true |
| species | *Toxoplasma gondii* ME49 | 1.0E-30 | false |
| genus | *Toxoplasma* | 1.0E-30 | false |
| species | *Plasmodium falciparum* | 1.0E-30 | false |

*merlin*-BIT models were generated on the "use specific BiGG models information" option, where we selected the template models.

*merlin* models will be referred to hereafter using the following nomenclature:

- *merlin*-DFA – *merlin* model using Diamond "fast" mode and Automatic workflow;
- *merlin*-DFS – *merlin* model using Diamond "fast" mode and *SamPler*;
- *merlin*-DSA – *merlin* model using Diamond "sensitive" mode and Automatic workflow;
- *merlin*-DSS – *merlin* model using Diamond "sensitive" mode and *SamPler*;
- *merlin*-BA – *merlin* model using BLAST and Automatic workflow;
- *merlin*-BS – *merlin* model using BLAST and *SamPler*.

### Results of models' assessment

#### *Bordetella pertussis*

#### Reactions

Seemingly, the best draft model obtained with *merlin* was generated with BIT. It achieved top performance on the most critical metrics: F1, Recall and JD. However, *merlin-DSA* and *merlin-DSS* obtained the best Precision and Ratio among *merlin* models. Besides *merlin-BIT* results, *merlin-BS* obtained the best F1 and JD among the non-template-based models. Nevertheless, the other *merlin* draft models achieved similar metric values, as the difference was on the order of the centesimal at most.

Table S5 – Reactions results for all the models of *B. pertussis*.

| model | Reactions number | Removed reactions | Converted reactions | Non-converted reactions | Duplicated reactions | recall | precision | f1 | TP | FP | FN | ratio | jaccard distance |
|---|---|---|---|---|---|---|---|---|---|---|---|---|---|
| ***merlin*-DFS** | 1891 | 601 | 607 | 683 | 6 | 0,31 | 0,47 | 0,37 | 282 | 319 | 642 | 0,88 | 0,77 |
| ***merlin*-DFA** | 1891 | 601 | 607 | 683 | 6 | 0,31 | 0,47 | 0,37 | 282 | 319 | 642 | 0,88 | 0,77 |
| ***merlin*-DSA** | 1761 | 570 | 560 | 631 | 6 | 0,29 | 0,49 | 0,37 | 272 | 282 | 652 | 0,96 | 0,77 |
| ***merlin*-DSS** | 1761 | 570 | 560 | 631 | 6 | 0,29 | 0,49 | 0,37 | 272 | 282 | 652 | 0,96 | 0,77 |
| ***merlin*-BS** | 1813 | 560 | 591 | 662 | 6 | 0,31 | 0,48 | 0,37 | 282 | 303 | 642 | 0,93 | 0,77 |
| **merlin-BA** | 1819 | 562 | 589 | 668 | 6 | 0,30 | 0,48 | 0,37 | 279 | 304 | 645 | 0,92 | 0,77 |
| ***merlin*-BIT** | 1526 | 337 | 1157 | 32 | 15 | 0,51 | 0,41 | 0,46 | 468 | 674 | 445 | 0,69 | 0,71 |
| **AuReMe** | 881 | 178 | 702 | 1 | 10 | 0,43 | 0,57 | 0,49 | 396 | 296 | 520 | 1,34 | 0,67 |
| **autoKEGGrec** | 884 | 0 | 575 | 309 | 7 | 0,34 | 0,55 | 0,42 | 310 | 258 | 614 | 1,20 | 0,74 |
| **CarveMe** | 2307 | 924 | 1375 | 8 | 63 | 0,53 | 0,37 | 0,43 | 482 | 830 | 426 | 0,58 | 0,72 |
| **ModelSEED** | 1336 | 185 | 554 | 597 | 1 | 0,29 | 0,49 | 0,37 | 273 | 280 | 660 | 0,98 | 0,77 |
| **PathwayTools** | 1334 | 35 | 379 | 920 | 1 | 0,25 | 0,62 | 0,36 | 235 | 143 | 695 | 1,64 | 0,78 |
| **RAVEN** | 3577 | 0 | 1133 | 2444 | 9 | 0,37 | 0,30 | 0,33 | 340 | 784 | 585 | 0,43 | 0,80 |

#### Genes

*merlin's* models performed similarly for gene sets, with no significant differences. However, it is worth noting that using *SamPler* seems to improve the annotation slightly, even if the sample annotation is performed automatically. Overall, the models generated with *merlin* outperformed almost all other models except for CarveMe in the most critical metrics.

Table S6 – Genes results for all the models of *B. pertussis*.

| model | Genes number | recall | precision | f1 | TP | FP | FN | ratio | jaccard distance |
|---|---|---|---|---|---|---|---|---|---|
| ***merlin*-DFS** | 832 | 0,66 | 0,61 | 0,63 | 507 | 325 | 263 | 1,56 | 0,54 |
| ***merlin*-DFA** | 829 | 0,66 | 0,61 | 0,63 | 507 | 322 | 263 | 1,57 | 0,54 |
| ***merlin*-DSA** | 738 | 0,63 | 0,66 | 0,65 | 487 | 251 | 283 | 1,94 | 0,52 |
| ***merlin*-DSS** | 737 | 0,63 | 0,66 | 0,65 | 487 | 250 | 283 | 1,95 | 0,52 |
| ***merlin*-BS** | 861 | 0,68 | 0,61 | 0,64 | 521 | 340 | 249 | 1,53 | 0,53 |
| **merlin-BA** | 858 | 0,67 | 0,60 | 0,64 | 519 | 339 | 251 | 1,53 | 0,53 |
| ***merlin*-BIT** | 814 | 0,65 | 0,61 | 0,63 | 497 | 317 | 273 | 1,57 | 0,54 |
| **AuReMe** | 452 | 0,47 | 0,80 | 0,59 | 362 | 90 | 408 | 4,02 | 0,58 |
| **autoKEGGrec** | 934 | 0,68 | 0,56 | 0,61 | 520 | 414 | 250 | 1,26 | 0,56 |
| **CarveMe** | 1026 | 0,76 | 0,57 | 0,65 | 584 | 442 | 186 | 1,32 | 0,52 |
| **ModelSEED** | 675 | 0,57 | 0,65 | 0,61 | 441 | 234 | 329 | 1,88 | 0,56 |
| **PathwayTools** | 780 | 0,60 | 0,60 | 0,60 | 465 | 315 | 305 | 1,48 | 0,57 |
| **RAVEN** | 1415 | 0,75 | 0,41 | 0,53 | 576 | 839 | 194 | 0,69 | 0,64 |

### *Lactobacilus plantarum*

#### Reactions

*merlin-BIT* outperformed the other models in all metrics except AuReMe and CarveMe (only in Recall). As for the other merlin models, *merlin-BS* was slightly better than *merlin-BA* for all metrics except Recall, while the other models surpassed *merlin-DFA*. Furthermore, it is worth noting that three out of the five top-performance models regarding F1, Ratio and JD were generated with *merlin*.

Table S7 – Reactions results for all the models of *L. plantarum*.

| model | Reactions number | Removed reactions | Converted reactions | Non-converted reactions | Duplicated reactions | recall | precision | f1 | TP | FP | FN | ratio | jaccard distance |
|---|---|---|---|---|---|---|---|---|---|---|---|---|---|
| ***merlin*-DFS** | 1201 | 320 | 427 | 454 | 6 | 0,43 | 0,49 | 0,46 | 205 | 216 | 274 | 0,95 | 0,71 |
| ***merlin*-DFA** | 1193 | 320 | 414 | 459 | 6 | 0,42 | 0,49 | 0,45 | 201 | 207 | 278 | 0,97 | 0,71 |
| ***merlin*-DSA** | 1205 | 324 | 432 | 449 | 6 | 0,43 | 0,48 | 0,46 | 206 | 220 | 273 | 0,94 | 0,71 |
| ***merlin*-DSS** | 1234 | 326 | 436 | 472 | 4 | 0,45 | 0,50 | 0,47 | 214 | 218 | 265 | 0,98 | 0,69 |
| ***merlin*-BS** | 1251 | 324 | 461 | 466 | 5 | 0,49 | 0,52 | 0,51 | 237 | 219 | 242 | 1,08 | 0,66 |
| **merlin-BA** | 1307 | 334 | 478 | 495 | 5 | 0,50 | 0,51 | 0,50 | 240 | 233 | 239 | 1,03 | 0,66 |
| ***merlin*-BIT** | 665 | 105 | 536 | 24 | 0 | 0,63 | 0,56 | 0,60 | 302 | 234 | 177 | 1,29 | 0,58 |
| **AuReMe** | 423 | 47 | 367 | 9 | 0 | 0,55 | 0,71 | 0,62 | 262 | 105 | 217 | 2,50 | 0,55 |
| **autoKEGGrec** | 673 | 0 | 441 | 232 | 7 | 0,49 | 0,54 | 0,51 | 235 | 199 | 244 | 1,18 | 0,65 |
| **CarveMe** | 1453 | 537 | 913 | 3 | 66 | 0,70 | 0,39 | 0,50 | 333 | 514 | 146 | 0,65 | 0,66 |
| **ModelSEED** | 1274 | 238 | 509 | 527 | 1 | 0,47 | 0,44 | 0,46 | 226 | 282 | 253 | 0,80 | 0,70 |
| **PathwayTools** | 2077 | 810 | 365 | 902 | 1 | 0,41 | 0,54 | 0,46 | 195 | 169 | 284 | 1,15 | 0,70 |
| **RAVEN** | 3184 | 0 | 1093 | 2091 | 9 | 0,58 | 0,26 | 0,36 | 278 | 806 | 201 | 0,34 | 0,78 |

#### Genes

*merlin-DFA* and *merlin-DFS* performed worse than the other drafts generated with *merlin*. These results suggest that Diamond's alignment results using the "fast" mode can affect the draft model construction while using the "sensitive" mode could not significantly improve or impair the reconstruction compared with BLAST. Furthermore, *merlin-BS* surpassed the other *merlin* models whose genome annotation was only performed with the automatic annotation algorithm (*merlin-BA, merlin-DFA*, and *merlin-DSA*), suggesting that *SamPler* can improve the similarity between a draft and a curated model.

Table S8 – Genes results for all the models of *L. plantarum*.

| model | Genes number | recall | precision | f1 | TP | FP | FN | ratio | jaccard distance |
|---|---|---|---|---|---|---|---|---|---|
| ***merlin*-DFS** | 606 | 0,61 | 0,73 | 0,67 | 444 | 162 | 284 | 2,74 | 0,50 |
| ***merlin*-DFA** | 595 | 0,60 | 0,73 | 0,66 | 435 | 160 | 293 | 2,72 | 0,51 |
| ***merlin*-DSA** | 635 | 0,64 | 0,73 | 0,68 | 464 | 171 | 264 | 2,71 | 0,48 |
| ***merlin*-DSS** | 631 | 0,63 | 0,72 | 0,67 | 456 | 175 | 272 | 2,61 | 0,50 |
| ***merlin*-BS** | 675 | 0,67 | 0,72 | 0,70 | 488 | 187 | 240 | 2,61 | 0,47 |
| **merlin-BA** | 725 | 0,67 | 0,68 | 0,68 | 491 | 234 | 237 | 2,10 | 0,49 |
| ***merlin*-BIT** | 552 | 0,61 | 0,81 | 0,70 | 445 | 107 | 283 | 4,16 | 0,47 |
| **AuReMe** | 317 | 0,40 | 0,92 | 0,56 | 291 | 26 | 437 | 11,19 | 0,61 |
| **autoKEGGrec** | 807 | 0,73 | 0,66 | 0,69 | 531 | 276 | 197 | 1,92 | 0,47 |
| **CarveMe** | 808 | 0,67 | 0,60 | 0,63 | 485 | 323 | 243 | 1,50 | 0,54 |
| **ModelSEED** | 718 | 0,67 | 0,68 | 0,67 | 488 | 230 | 240 | 2,12 | 0,49 |
| **PathwayTools** | 856 | 0,70 | 0,59 | 0,64 | 507 | 349 | 221 | 1,45 | 0,53 |
| **RAVEN** | 1295 | 0,84 | 0,47 | 0,60 | 609 | 686 | 119 | 0,89 | 0,57 |

***Toxoplasma gondii***

### Reactions

*merlin-DFS, -DSA,* and *merlin-DSS* models presented neglectable differences for all metrics, which indicate they are extremely similar. On the other hand, *merlin-BIT* obtained the worst results overall except for Precision and Ratio.

Table S9 – Reactions results for all the models of *T. gondii*.

| model | Reactions number | Removed reactions | Converted reactions | Non-converted reactions | Duplicated reactions | recall | precision | f1 | TP | FP | FN | ratio | jaccard distance |
|---|---|---|---|---|---|---|---|---|---|---|---|---|---|
| ***merlin*-DFS** | 1345 | 215 | 1125 | 5 | 8 | 0,61 | 0,51 | 0,55 | 570 | 547 | 370 | 1,04 | 0,62 |
| ***merlin*-DFA** | 1446 | 228 | 1211 | 7 | 8 | 0,63 | 0,49 | 0,55 | 593 | 610 | 347 | 0,97 | 0,62 |
| ***merlin*-DSA** | 1335 | 210 | 1120 | 5 | 8 | 0,60 | 0,51 | 0,55 | 567 | 545 | 373 | 1,04 | 0,62 |
| ***merlin*-DSS** | 1335 | 210 | 1120 | 5 | 8 | 0,60 | 0,51 | 0,55 | 567 | 545 | 373 | 1,04 | 0,62 |
| ***merlin*-BS** | 1441 | 228 | 1206 | 7 | 7 | 0,63 | 0,49 | 0,55 | 591 | 608 | 349 | 0,97 | 0,62 |
| **merlin-BA** | 1441 | 228 | 1206 | 7 | 7 | 0,63 | 0,49 | 0,55 | 591 | 608 | 349 | 0,97 | 0,62 |
| ***merlin*-BIT** | 564 | 106 | 134 | 324 | 9 | 0,09 | 0,71 | 0,17 | 89 | 36 | 853 | 2,47 | 0,91 |
| **AuReMe** | 432 | 89 | 209 | 134 | 7 | 0,18 | 0,83 | 0,29 | 167 | 35 | 774 | 4,77 | 0,83 |
| **autoKEGGrec** | 567 | 0 | 564 | 3 | 7 | 0,43 | 0,72 | 0,53 | 400 | 157 | 541 | 2,55 | 0,64 |
| **PathwayTools** | 1533 | 243 | 356 | 934 | 2 | 0,26 | 0,69 | 0,38 | 245 | 109 | 695 | 2,25 | 0,77 |
| **RAVEN** | 2655 | 0 | 2640 | 15 | 14 | 0,90 | 0,32 | 0,48 | 847 | 1779 | 93 | 0,48 | 0,69 |

### Genes

Apart from *merlin-DSA* and *-DSS* outperforming the other merlin models, their differences were negligible. Moreover, using *SamPler* was not as relevant as for the prokaryote reconstructions.

Table S10 – Genes results for all the models of *T. gondii*.

| model | Genes number | recall | precision | f1 | TP | FP | FN | ratio | jaccard distance |
|---|---|---|---|---|---|---|---|---|---|
| ***merlin*-DFS** | 591 | 0,60 | 0,53 | 0,56 | 315 | 276 | 211 | 1,14 | 0,61 |
| ***merlin*-DFA** | 715 | 0,65 | 0,47 | 0,54 | 333 | 382 | 181 | 0,87 | 0,63 |
| ***merlin*-DSA** | 589 | 0,60 | 0,53 | 0,57 | 315 | 274 | 210 | 1,15 | 0,61 |
| ***merlin*-DSS** | 589 | 0,60 | 0,53 | 0,57 | 315 | 274 | 210 | 1,15 | 0,61 |
| ***merlin*-BS** | 708 | 0,63 | 0,46 | 0,53 | 324 | 384 | 193 | 0,84 | 0,64 |
| **merlin-BA** | 708 | 0,63 | 0,46 | 0,53 | 324 | 384 | 193 | 0,84 | 0,64 |
| ***merlin*-BIT** | 104 | 0,07 | 0,62 | 0,13 | 64 | 40 | 794 | 1,60 | 0,93 |
| **AuReMe** | 243 | 0,20 | 0,62 | 0,31 | 150 | 93 | 587 | 1,61 | 0,82 |
| **autoKEGGrec** | 772 | 0,51 | 0,38 | 0,43 | 293 | 479 | 287 | 0,61 | 0,72 |
| **PathwayTools** | 1236 | 0,72 | 0,28 | 0,40 | 345 | 891 | 133 | 0,39 | 0,75 |
| **RAVEN** | 1017 | 0,93 | 0,38 | 0,54 | 388 | 629 | 30 | 0,62 | 0,63 |
