## Supplementary data 3 for "*merlin* v4.0: an updated platform for the reconstruction of high-quality genome-scale metabolic models"

* To whom correspondence should be addressed.

### Supplementary data 3 – *merlin's* graphical interface

*merlin's* graphical interface has changed considerably since the last publication. The main views are shown in the following figures.

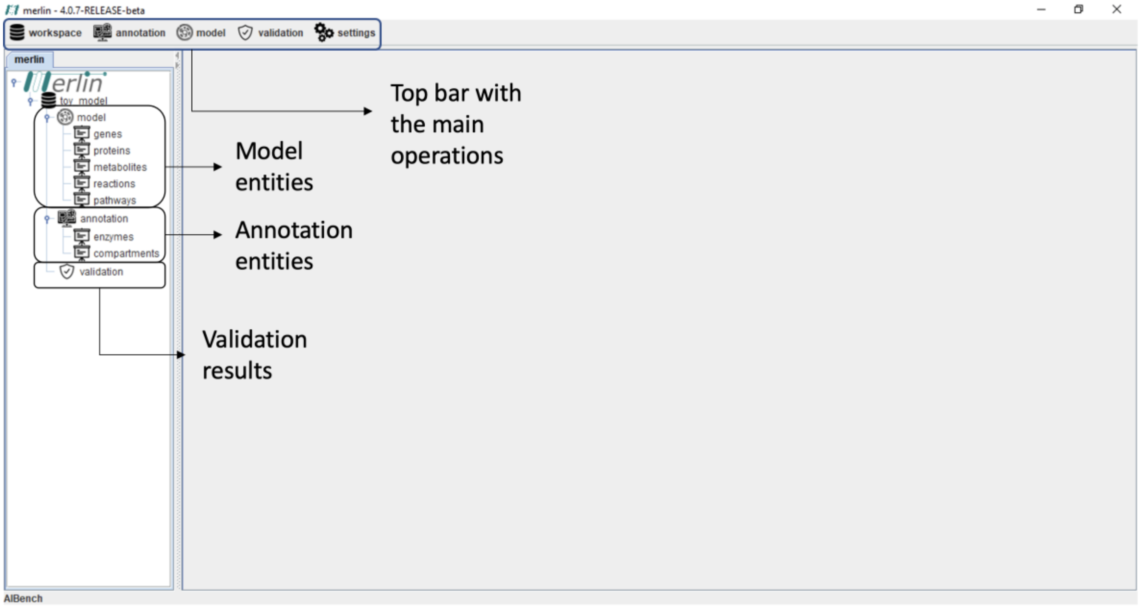


**Figure S1 –** Main view and entities of *merlin*. At the top bar, several menus associated with the primary operations are rendered. These operations are associated with the workspaces' management, annotations, model manipulation and refinement, validation and curation, settings for defining several parameters, installing or updating plugins, and others. Moreover, the model, annotation, and validation entities are associated with each workspace in the dashboard on the left. The model entity includes *genes, proteins, metabolites, reactions, and pathways* subentities; Whereas the annotation entity includes *compartments* and *enzymes* subentities; finally, the validation renders the results of the BioISO analysis.

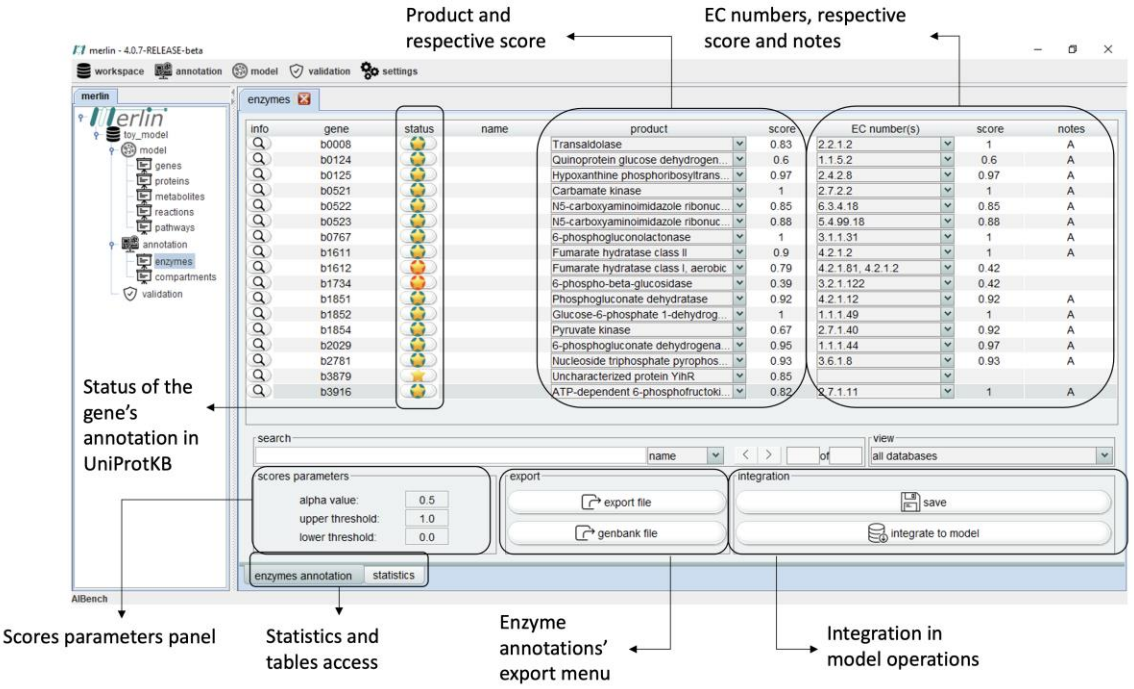


**Figure S2 –** Enzymes' view in *merlin*'s graphical interface. It contains information regarding the enzymes annotation. Moreover, several features are here provided, namely: the Enzyme Commission (EC) number and products' scores, the gene's annotation status and notes written by the user or by annotation tools such as *SamPleR* or *Automatic Workflow*. The view helps in manual curation, and all alterations can be integrated into the model at any time.

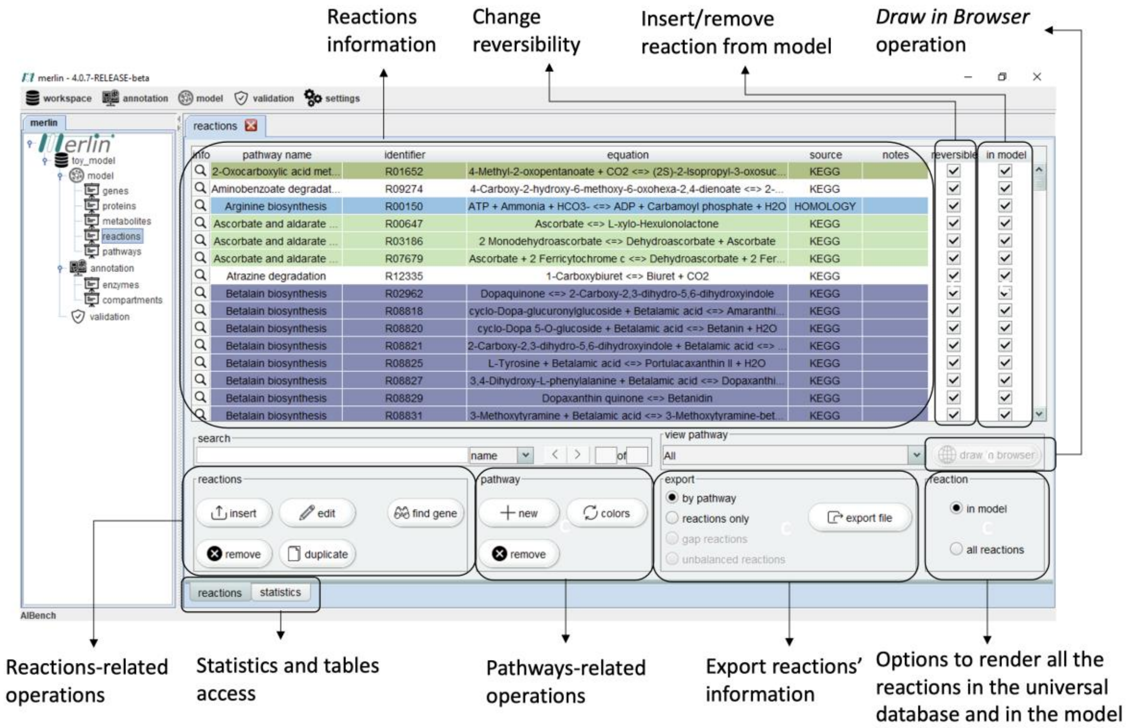


**Figure S3 –** Reactions' view in *merlin*'s graphical interface. All the reactions in the model are enumerated in the main table. Moreover, the universal database of reactions is shown when the respective option is selected (bottom right corner). Notwithstanding, several operations such as reactions insertion, edition, duplication and deletion are here provided. Furthermore, pathways can be added, removed and visualised using the *Draw in Browser* operation on the right.

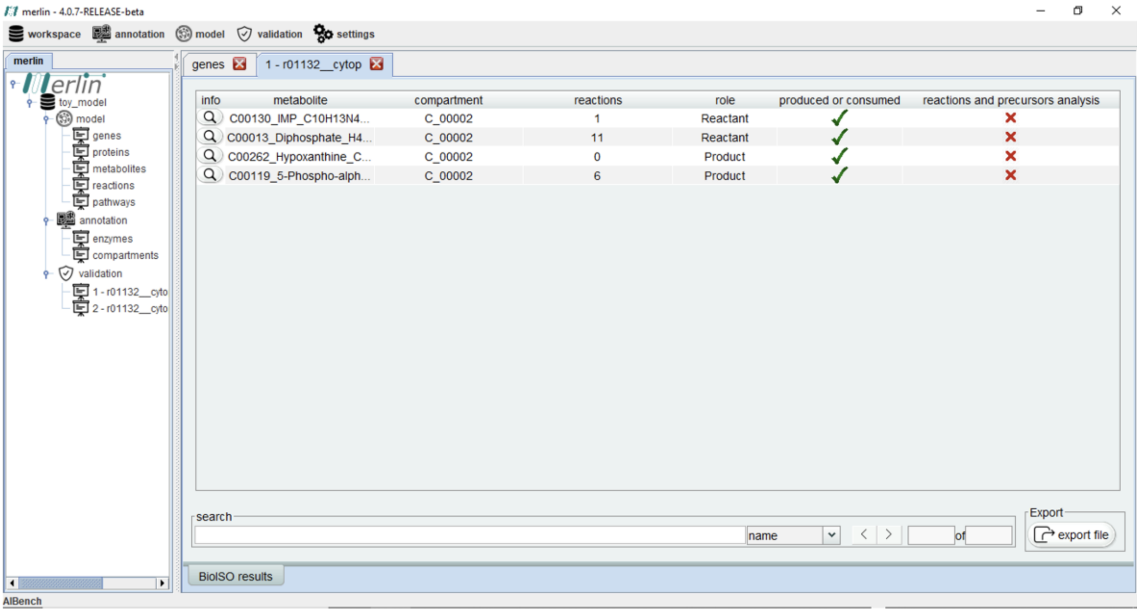


**Figure S4 –** BioISO's results view in *merlin*'s graphical interface.

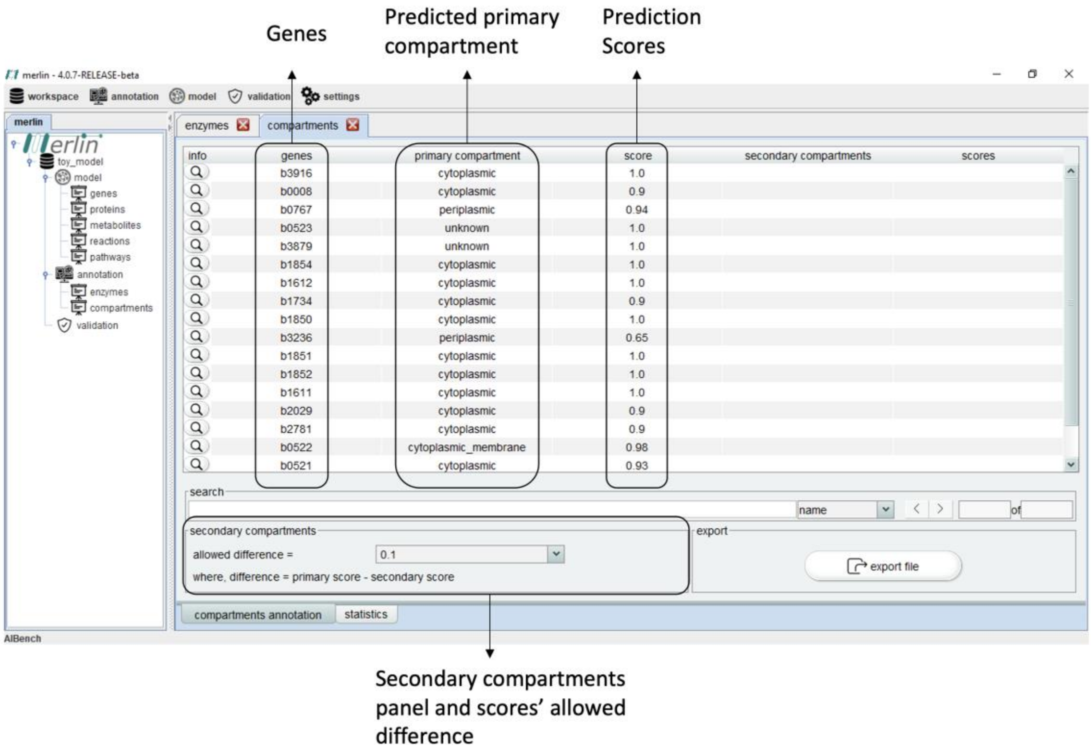


**Figure S5 –** Compartments' view in *merlin*'s graphical interface. It provides the localisation of the enzymes encoded in the genome associated with the model's reaction. Scores for the localisation prediction are shown, and potential secondary compartments may be included, depending on the tool. The user can change the acceptable difference between the primary and secondary compartment score in the dropdown box in the secondary compartments panel. If the scores' difference is lower than the one previously set, the reaction catalysed by the encoded protein will occur in two compartments.

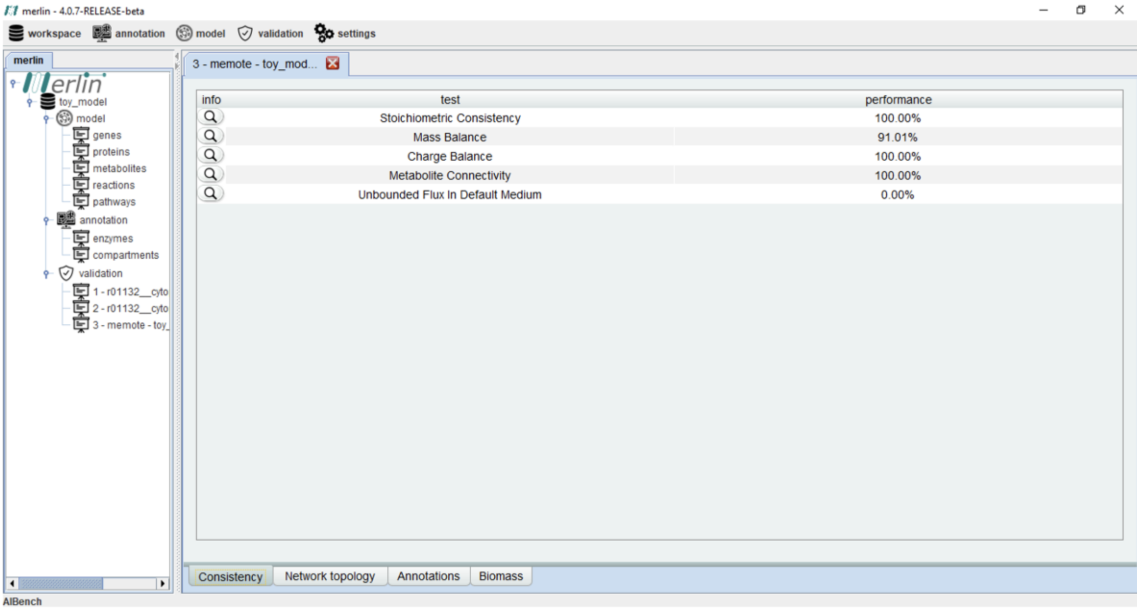


**Figure S6 –** MEMOTE results' view in *merlin*'s graphical interface.
